# Accounting for age-of-onset and family history improves power in genome-wide association studies

**DOI:** 10.1101/2021.04.20.440585

**Authors:** Emil M Pedersen, Esben Agerbo, Oleguer Plana-Ripoll, Jakob Grove, Julie W. Dreier, Katherine L. Musliner, Marie Bækvad-Hansen, Georgios Athanasiadis, Andrew Schork, Jonas Bybjerg-Grauholm, David M. Hougaard, Thomas Werge, Merete Nordentoft, Ole Mors, Søren Dalsgaard, Jakob Christensen, Anders D. Børglum, Preben B. Mortensen, John J. McGrath, Florian Privé, Bjarni J. Vilhjálmsson

## Abstract

Genome-wide association studies (GWAS) have revolutionized human genetics, allowing researchers to identify thousands of disease-related genes and possible drug targets. However, case-control status does not account for the fact that not all controls may have lived through their period of risk for the disorder of interest. This can be quantified by examining the age-of-onset distribution and the age of the controls or the age-of-onset for cases. The age-of-onset distribution may also depend on information such as sex and birth year. In addition, family history is not routinely included in the assessment of control status. Here we present LT-FH++, an extension of the liability threshold model conditioned on family history (LT-FH), that jointly accounts for age-of-onset and sex, as well as family history. Using simulations, we show that, when family history and the age-of-onset distribution are available, the proposed approach yields large power gains over both LT-FH and genome-wide association study by proxy (GWAX). We applied our method to four psychiatric disorders available in the iPSYCH data, and to mortality in the UK Biobank, finding 20 genome-wide significant associations with LT-FH++, compared to 10 for LT-FH and 8 for a standard case-control GWAS. As more genetic data with linked electronic health records become available to researchers, we expect methods that account for additional health information, such as LT-FH++, to become even more beneficial.

## Introduction

Identifying the genetic variants underlying diseases and traits is a hallmark of human genetics. In recent years, large meta-analyses of genome-wide association studies (GWAS) have identified thousands of genetic variants for common diseases^1–7^ including psychiatric disorders^8–12^, revealing a remarkably complex and polygenic genetic architecture for most traits. International research collaboration where GWAS summary statistics have been shared in large consortia has been vital to this success, allowing researchers to obtain large sample sizes needed to study polygenic diseases. Novel advances in computational methods have also contributed to this success by enabling researchers to do more with less data^13–17^. Yet, for most of these traits and diseases only a small fraction of the estimated heritable variation has been identified in GWAS^18,19^, highlighting the need for even larger samples and more powerful analysis methods.

Currently, most case-control GWAS studies are conducted using a regression model where the outcome is the case-control status, or occasionally, the age-of-onset of disease^20^. In this paper, we have opted for using the phrase age-of-onset over age-at-first-diagnosis, since they commonly refer to the same underlying thing, i.e. when a diagnosis is given. Recently, researchers have proposed several methods that leverage additional information to improve the power to detect genetic associations, without having to increase the number of genotyped individuals. These include multivariate methods that leverage shared environmental or genetic correlations between traits and diseases^21–25^, as well as methods that account for age-of-onset^26,27^. Perhaps the most fruitful development has come from methods that leverage family information to increase statistical power to identify associations, such as genome-wide association study by proxy (GWAX)^28,29^ and liability threshold model based approach^30^. The liability threshold model conditioned on family history (LT-FH)^30^ estimates the posterior mean genetic liability under the liability threshold model conditional on the case-control status of the individual, parents, and siblings. Here, *family history* refers to the case-control status of all family members, i.e. parents and siblings. As for GWAX, it considers any individual with a family member, who has the disorder being studied, as a case, increasing the number of cases. The GWAX phenotype remains a case-control phenotype. Although both GWAX and LT-FH can lead to power increases over case-control GWAS on real data, they achieve it in two different ways. It has been shown that GWAX can lead to a reduction in power when compared to a case-control GWAS, if the in-sample disease prevalence is high. However, LT-FH consistently provides an increase in power compared to case-control GWAS and GWAX^30^. This power improvement in LT-FH stems from two main sources. First, it distils family information and the individual’s case-control status into a genetic liability estimate, resulting in a more informative outcome than the case-control status alone, to be used in GWAS. Second, it also allows researchers to include more individuals in their analysis. For instance when studying breast cancer we can derive the posterior genetic liability for genotyped males conditional on the family history for their mothers and sisters, and thus include them in the GWAS.

However, family members often span a large age-range, which can affect the expected disease prevalence due to changes in diagnostic methods and criteria over time. We refer to such differences in prevalence by birth year as *cohort effects*. For instance, in the iPSYCH data^31^, where genotyped individuals are born after 1980, we expect severe right censoring for many diagnoses. Survival models are routinely used in epidemiology to model time-to-event data in order to account for right censoring, time at risk, and age-of-onset, as well as cohort effects^32^. They have also been shown to provide up to 10% increase in power to detect genetic variants in GWAS when compared to standard logistic regression^26^. More recently, computationally efficient random effect survival models (frailty models) that can control for population and family structure have been proposed for GWAS^27^. However, although powerful, survival model GWAS methods that account for family history as well have yet to be proposed. The survival model is based on a fundamentally different generative model than the liability threshold model and the posterior genetic liabilities derived with LT-FH cannot be used directly as an outcome in a survival analysis. Hujoel *et al.^30^* proposed an approach to address this problem by accounting for age-of-onset in the genotyped individuals by linearly shifting the threshold for the genetic liabilities based on observed in-sample prevalence in different age groups, but did not observe any improvements in power. We believe that this approach was unsuccessful in part because the in-sample estimate of the prevalence is subject to both a survival and selection bias, and does not properly reflect prevalence in the population.

In this paper we propose LT-FH++, a method that extends the model underlying LT-FH to account for information such as right censoring, age-of-onset, sex, and cohort effects. We achieve this by using a personalized threshold for each person (including family members), conditional on available information as well as general population incidence rates by age, sex, and birth year. LT-FH++ has been implemented into an R package (See Code Availability), which utilizes a Gibbs sampler implemented in C++ through the Rcpp R package^33^. The personalized thresholds are made possible by replacing the Monte Carlo sampling used by Hujoel et al. with a much more efficient Gibbs sampler. The Gibbs sampler allows us to estimate the posterior mean genetic liability for each individual independent of one another, thereby making it highly scalable.

First, we perform a GWAS with the standard case-control phenotype as well as GWAX, LT-FH, and LT-FH++ outcomes for simulated data with the liability threshold model as the generative model. For real-world application, we analysed mortality in the UK biobank and 4 psychiatric disorders in the iPSYCH cohort. Of the 4 psychiatric disorders, we will specifically highlight ADHD, since it had the highest number of genome-wide significant hits.

## Results

### Overview of methods

The LT-FH++ method proposed here extends the LT-FH method to account for additional information for family members, such as age, sex, and cohort effects for case-control outcomes. LT-FH assumes a liability threshold model, where every individual has an underlying liability for the outcome, but only becomes a case if the liability exceeds a given threshold, which is determined by the sample or population prevalence^34^. It further assumes that the covariance structure depends on the heritability and relatedness coefficient between each individual, which is a reasonable assumption for polygenic case-control diseases^35,36^. Under these assumptions, LT-FH estimates the posterior mean genetic liability conditional on the case-control status of the genotyped individual and their family members using a Monte Carlo sampling. The posterior mean genetic liability is then used as the continuous outcome in a GWAS, e.g. using BOLT-LMM^37^.

In LT-FH++ we introduce an *age-dependent liability threshold model* to capture the effect of age, and replace the Monte Carlo sampling with a much more computationally efficient Gibbs sampler. Illustrated in **Figure 1A**, the age-dependent liability threshold model extends the liability threshold model by assuming that the threshold for becoming a case at a given age corresponds to the prevalence of the disease at that age. Interestingly, this model can be viewed as a type of survival analysis (see methods). We can then account for additional information, such as birth year and sex, by further conditioning the disase prevalence on this information. This leads to an individualized disease liability threshold for each person, including family members, which in practice requires us to be able to estimate separate genetic liabilities for each individual. This is made possible by replacing the Monte Carlo strategy of LT-FH with the computationally efficient Gibbs sampler that can sample from multivariate truncated Gaussian distributions to obtain personalized genetic liability estimates. As illustrated in **Figure 1B** this results in more precise genetic liability estimates for LT-FH++ under the model compared to LT-FH, which for a population translates also into more variable genetic liability estimates (see **Figure S1**). Thus, in order to reap the full benefit of LT-FH++ it requires prevalence information to be available by age, sex, and birth year. Fortunately, such information is often partially or fully available on a population-level, e.g. in the Danish registers^38^. Using population prevalence information also allows LT-FH++ to estimate the genetic liability on a population-scale, which may also reduce the risk of ascertainment and selection bias^39–41^. We summarize the information that LT-FH++ can account for and the two step procedure of estimating individual genetic liabilities and performing GWAS on these in **Figure 1C**.

**Figure 1:**
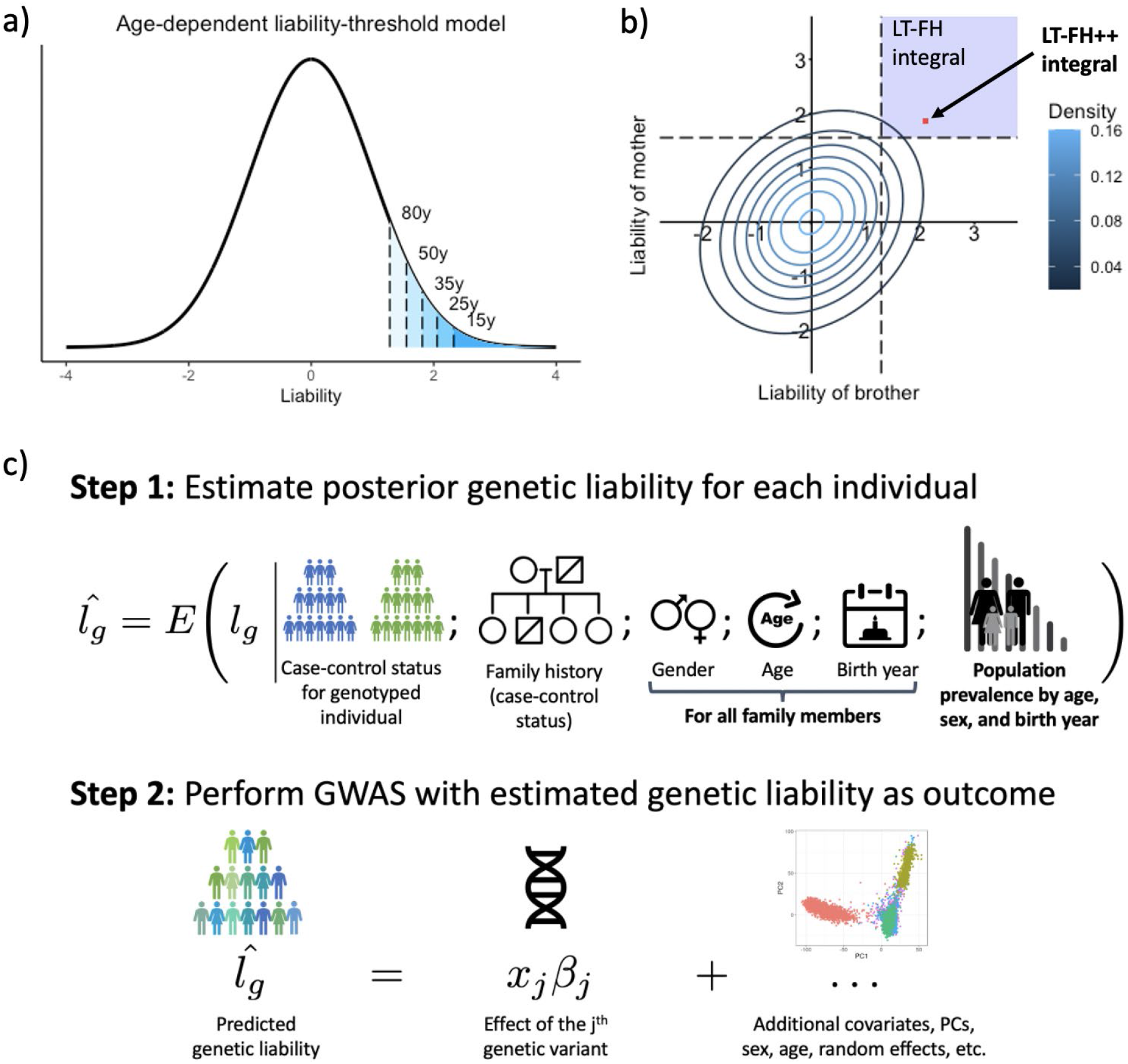
Illustration of the differences between LT-FH and LT-FH++. In **A)**, an age-dependent liability threshold model with different thresholds marked. The marks correspond to the prevalence at the age of 80y (10%), 50y (6%), 35y (3.5%), 25y (2%), and 15y (1%). **B)** The posterior mean estimate of the liability is obtained by integrating over the liability space spanned by the genotyped individual and its family members. Here, we consider a brother and a mother, where the contour lines indicate the joint multivariate liability density of the mother and the brother (assuming a heritability of 0.5). Using fixed population prevalence for males and females (dashed lines), and assuming mother and brother are cases, LT-FH integrates over the blue shaded area to estimate the genetic liability. In contrast LT-FH++ considers the age-of-onset, sex, and birth year for family members to obtain a more precise genetic liability estimate highlighted by the red dot. **C)** An overview of how LT-FH++ GWAS works, and what information it accounts for. In contrast to LT-FH, which accounts for the case-control status of the genotyped individual and family history, LT-FH++ also uses population prevalence information to account for gender, age, and birth year of family members. As with LT-FH, the predicted liabilities are then used as a continuous outcome in a GWAS using BOLT-LMM^37^.

### Simulation Results

We examined the performance of LT-FH++ using both simulated and real data. We simulated 100,000 unrelated individuals each with 100,000 independent single-nucleotide polymorphisms (SNPs), and their family (two parents and 0-2 siblings). We generated case-control outcomes under the liability threshold model, and assigned age-of-onset by assuming the prevalence followed a logistic curve as a function of age (see Methods section for simulation details).

We first considered the simulations for families with no siblings. We benchmarked LT-FH++ against case-control status and LT-FH. The results for 5% prevalence are shown in **Figure 2**, and the results for 10% prevalence can be found in **Figure S2**. We simulated sample ascertainment by downsampling controls such that cases and controls had equal proportions (50% each), which translated into a total of 10.000 individuals for a 5% prevalence and 20.000 individuals for a 10% prevalence. The simulation results confirmed the increase in power (number of causal SNPs detected) of LT-FH over standard GWAS when accounting for family history^30^. When also accounting for sex differences and age in LT-FH++, we observed a further increase in power, especially when the cases were ascertained (downsampling controls). Averaging over 10 simulations, LT-FH had a power improvement over standard GWAS between 13.9% and 54.4%, where less power improvement was observed when downsampling controls. In contrast, the average power increase for LT-FH++ and standard GWAS was between 34.2% and 60.6%. Without downsampling controls, the relative improvements of LT-FH++ over LT-FH for a 5% and 10% prevalence were 3.98% and 4.81%, respectively. However, when downsampling controls, we observed an improvement of 17.8% for a 5% prevalence and 14.8% for a 10% prevalence. Similar results are obtained when simulating two siblings in families (results not shown).

**Figure 2:**
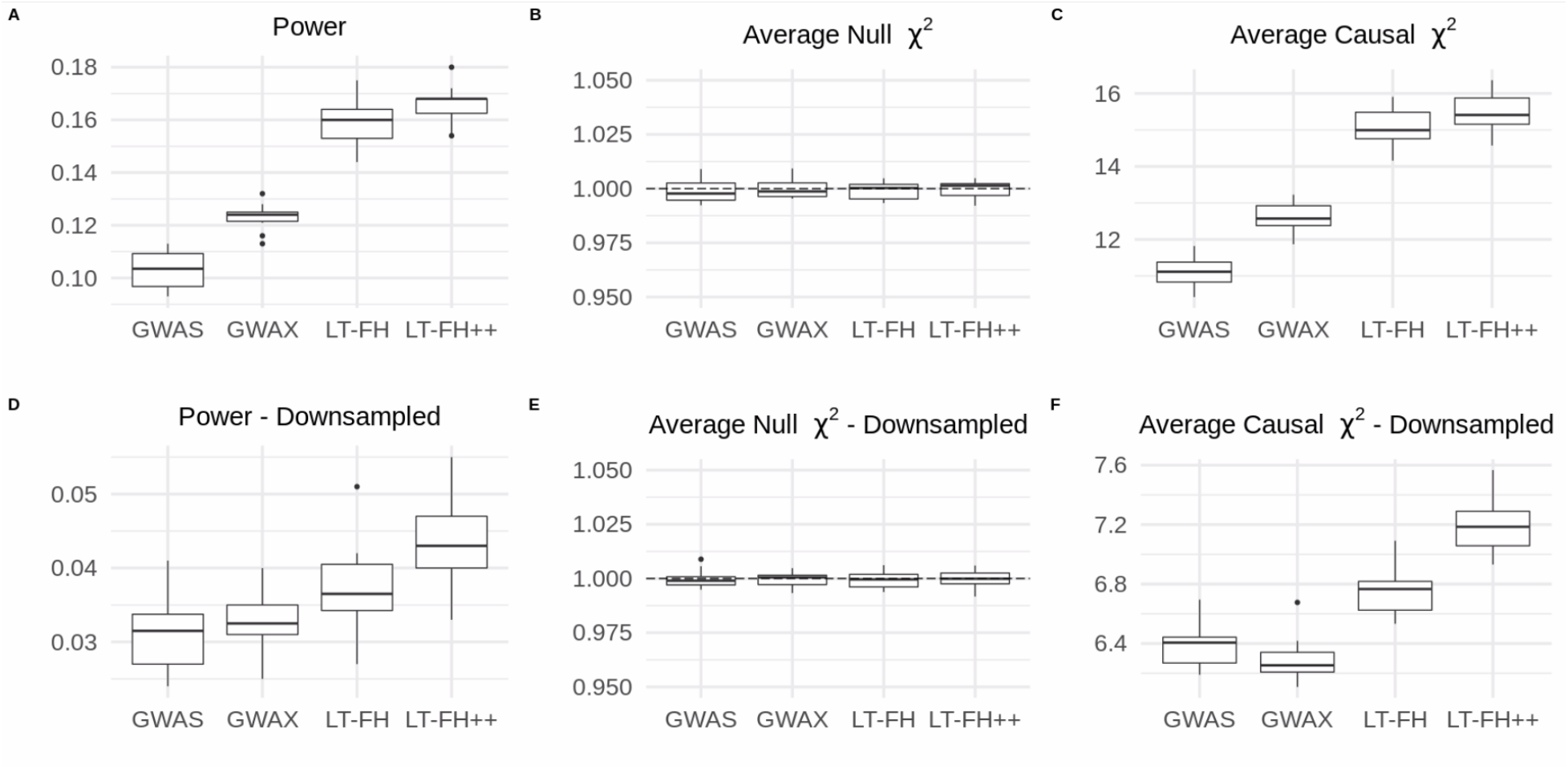
Simulation results for a 5% prevalence, with and without downsampling of controls. Linear regression was used to perform the GWAS for LT-FH and LT-FH++, while a 1-df chisq test was used for case-control status. We assessed the power of each method by considering the fraction of causal SNPs with a p-value below 5 × 10^−8^. Here GWAS refers to case-control status and LT-FH and LT-FH++ are both without siblings. Downsampling refers to downsampling the controls such that we have equal proportions of cases and controls, i.e. we have 10,000 individuals total for a 5% prevalence and 20,000 individuals for a 10% prevalence.

We also assessed the robustness of LT-FH++ by misspecifying model hyper-parameters, i.e. the heritability and prevalence parameters. Simulated heritability was 50% and when misspecifying it we used 25% and 75%. For the prevalence, we used simulated values of either 5% or 10%, and used either half or double of the true value to assess the impact of misspecifying this parameter. This resulted in e.g. a prevalence of 5% or 20% when the true prevalence was 10%. In **Figures S3-6**, when misspecifying the heritability and prevalence, we see similar results as in **Figure 2** with nearly identical mean null *χ*^2^ statistics, mean causal *χ*^2^ statistics, and power. LT-FH++ is therefore robust to misspecification of heritability and prevalence.

### Analysis of mortality in the UK biobank

To evaluate the performance of LT-FH++ on real data we chose mortality in the UK biobank, as this is the only outcome available where we have age information for family members, i.e. we have age or age-of-death for mothers and fathers. We then obtained population prevalence information from the Office for National Statistics (ONS), which provides mortality rates for England and Wales by sex and birth year (since 1841), and for the United Kingdom (UK) since 1950. This allowed us to obtain individualized prevalence thresholds for LT-FH++ for each genotyped individual and their parents (see Methods for details). The mortality rates by age and sex are shown for each decade in **Figure S7**.

The Manhattan plots for standard case-control, LT-FH, and LT-FH++ GWAS, can be found in **Figure 3** (see Methods for analysis details). When using the case-control phenotype as the outcome in GWAS, we did not observe any genome-wide significant SNPs. For LT-FH, we found two genome-wide significant loci, including a well known association with mortality in the *APOE* gene^42^ and in the *HYKK* gene, which is strongly associated with smoking behavior^6^. These were also the two strongest associations found with LT-FH++, which additionally found 8 other independent associated variants, where independence was assessed using GCTA-COJO^43^. The 10 identified variants are shown in **Table S1**, of which three variants have not previously been identified as associated with mortality or aging. One of these is near the *HLA-B* gene, which is involved in immune response and has been found to be associated with white blood cell count^44^ and Psoriasis^45^. The second association is near the *MYCBP2* gene, which has previously been identified as being associated with chronotype^46^, and the expression of this gene was recently found to increase with age and interact with the SARS-CoV-2 proteome^47^. The third association was near the *ZBBX* gene, which has been found to be associated with changes in DNA methylation with age^48^.

**Figure 3:**
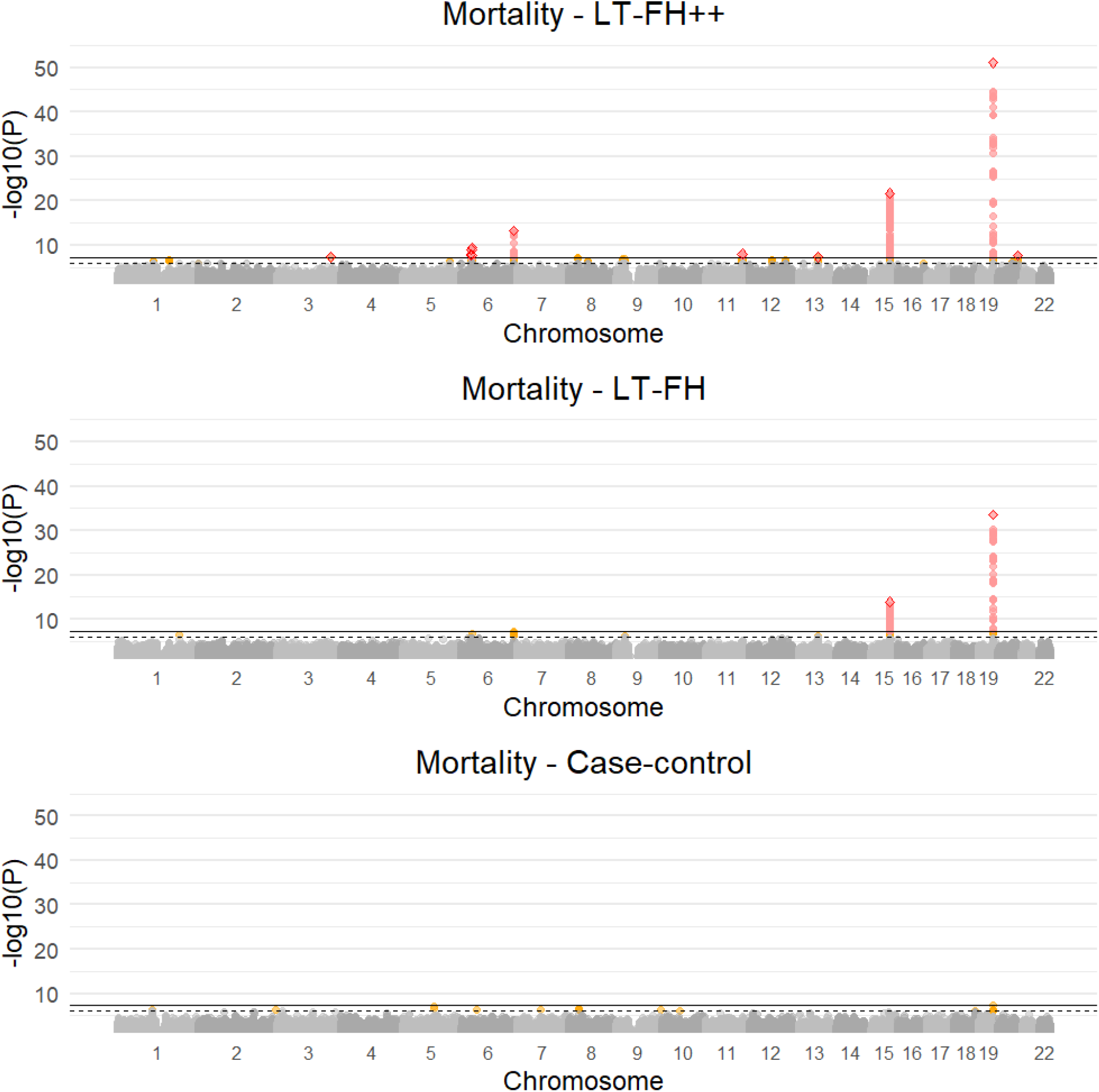
Manhattan plots for LT-FH++, LT-FH, and case-control GWAS of mortality in the UK Biobank. The Manhattan plots display a Bonferroni corrected significance level of 5 × 10^−8^, and a suggestive threshold of 5 × 10^−6^. The genome-wide significant SNPs are colored in red and the suggestive SNPs are colored in orange. The squares correspond to top SNPs in a window of size 300k base pairs.

The power increase between two GWAS outcomes can be assessed by plotting the Z-scores against each other. For LT-FH++, it leads to an estimated power increase of 49% over LT-FH. Since the Z-scores squared are the *χ*^2^ statistics, we opted to illustrate the power improvement of LT-FH++ over LT-FH, through the *χ*^2^ statistics. We plotted the *χ*^2^ statistic for variants with a p-value below 5 × 10^−6^ in **Figure 4**. LT-FH and LT-FH++ both had a large increase in power over case-control status, resulting in an estimated relative power increase of 75% and 187%, respectively. The *χ*^2^ statistics and Z-scores plots compared to case-control status can be found in **Figures S8 and S9**.

**Figure 4:**
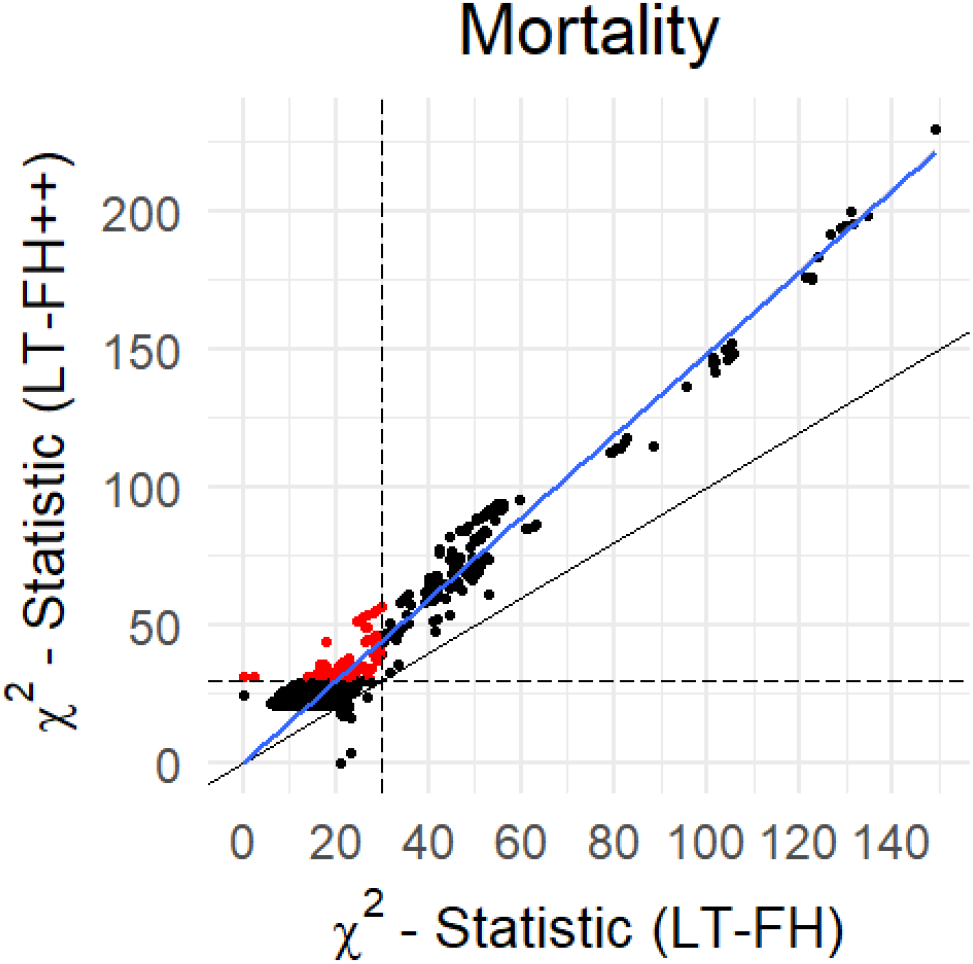
The *χ*^2^ statistics for LT-FH++ versus the ones for LT-FH for the GWAS of mortality in the UK Biobank. We restricted to variants with a p-value below 5 × 10^−6^ for at least one of the compared outcomes. The red dots are variants identified as genome-wide significantly associated by only one of the outcomes. The black dots are suggestive associations identified by either method, or genome-wide significant associations identified by both methods. The black line indicates the identity line and the blue line is the best fitted line using linear regression. The black dashed lines correspond to the threshold for genome-wide significance.

### Application to four psychiatric disorders in iPSYCH

The iPSYCH data^31^ with linked Danish registers has age and age-of-onset information for all close family members of genotyped individuals. We considered four psychiatric disorders in the iPSYCH data, ADHD, Autism Spectrum Disorder (ASD), Depression, and Schizophrenia. For each of these we obtained prevalences by birth-year, age, and sex using the same diagnostic criteria (see Methods for details). As shown in **Figures S10-13** the prevalence of psychiatric disorders strongly depend on birth year and sex, making it an appealing application of LT-FH++. We performed a GWAS of the three outcomes, case-control GWAS, LT-FH, and LT-FH++ for the four psychiatric disorders (see Methods for analysis details). Across the four psychiatric disorders we found 10 genome-wide significant variants using LT-FH++ compared to 8 for LT-FH and case-control. For ADHD, LT-FH++ found 7 genome-wide significant loci, he two additional identified variants were for ADHD, on chromosome 11 near the *LINC02758* gene which was found to be associated with ADHD in a meta-analysis^10^, and another on chromosome 14 in the *AKAP6* gene, which has previously been identified as being associated with cognitive traits^49,50^. The Manhattan plots for ADHD can be seen in **Figure 5** for all three outcomes, i.e. case-control, LT-FH and LT-FH++ (see Methods for details). Manhattan plots for all three outcomes are very similar, with no one outcome clearly outperforming the others. However, LT-FH++ does have two genome-wide significant SNPs that were close to genome-wide significance with both LT-FH and case-control analysis, but did not pass the significance threshold. Similarly, LT-FH++ and case-control have one SNP that is not found by LT-FH, however it is also close to the genome-wide significance threshold for LT-FH. In **Figure 6** we show the *χ*^2^ statistics plot restricting to LD clumped SNPs with a p-value threshold of 5 × 10^−6^ for the index SNP and the clumped SNPs from the largest external meta analysed ADHD summary statistics (see Methods for details). If one method had clearly performed better than another, we would have expected to see a slope different from one, however this is not the case here. Overall, there is little power improvement by using either LT-FH or LT-FH++ over case-control GWAS for ADHD.

**Figure 5:**
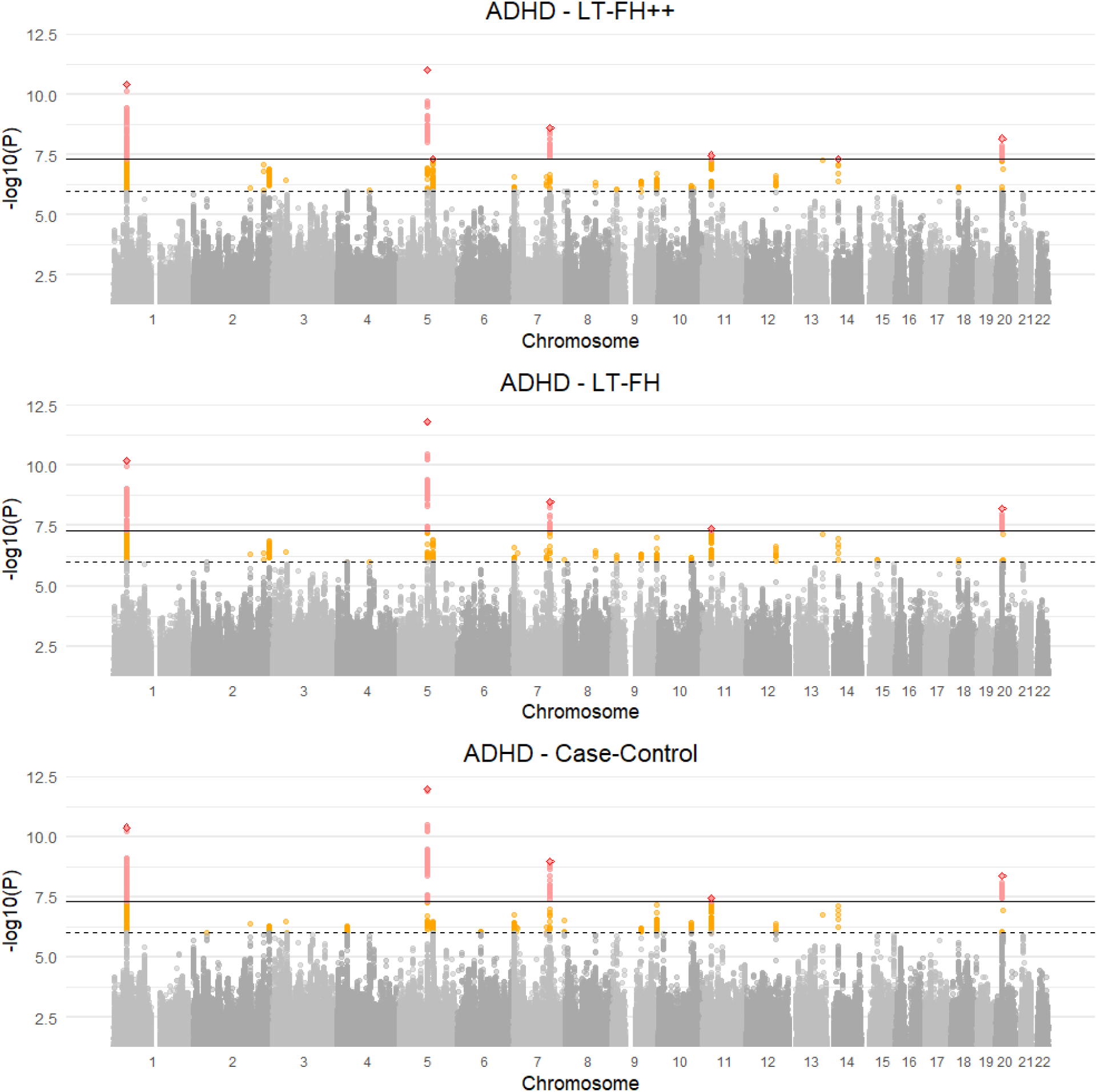
Manhattan plots for LT-FH++, LT-FH, and case-control GWAS of ADHD in the iPSYCH data. The dashed line indicate a suggestive p-value of 5 × 10^−6^ and the fully drawn line at 5 × 10^−8^ indicates genome-wide significance threshold. The suggestive SNPs are colored in orange, while the genome-wide significant SNPs are colored in red. The squares correspond to top SNPs in a window of size 300k base pairs.

**Figure 6:**
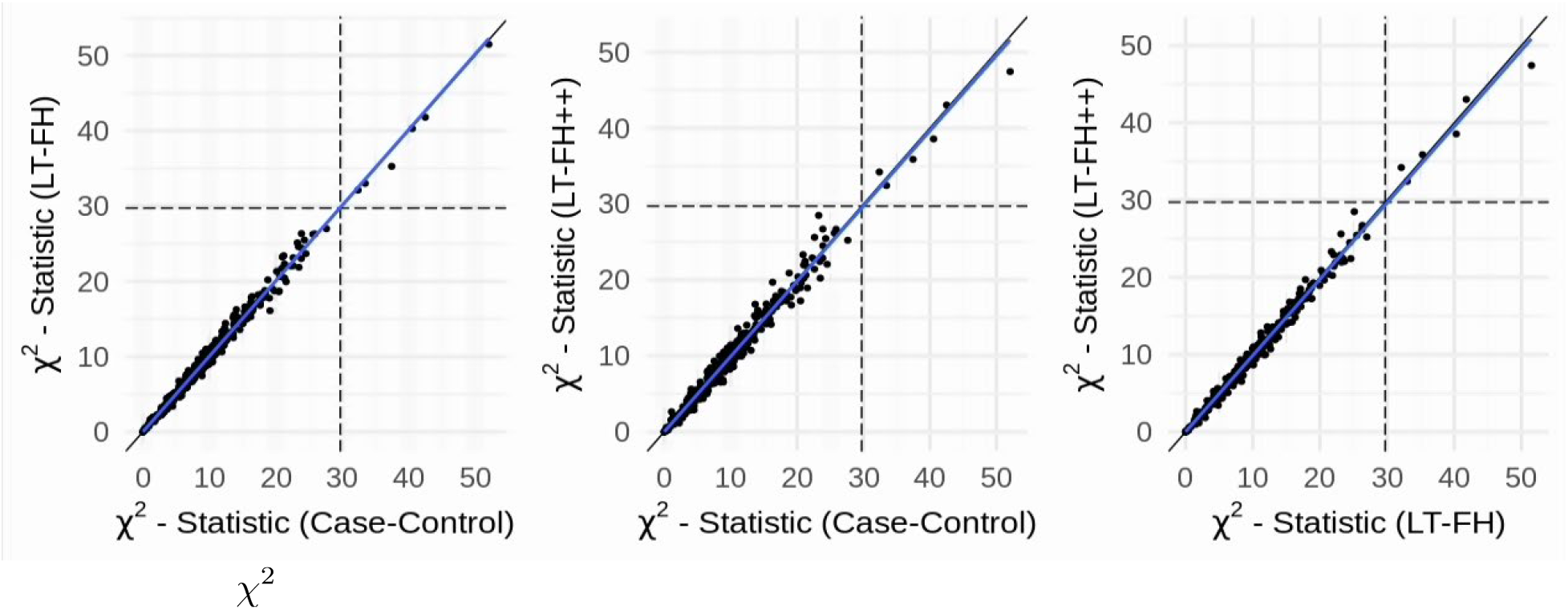
The *χ*^2^ statistics from the GWAS of ADHD for each of the three methods (LT-FH++, LT-FH, and case-control GWAS), plotted against each other. The dots correspond to LD clumped SNPs that have a p-value below 5 × 10^−6^ in the largest published meta analysis and present in the iPSYCH cohort (see Methods for details). The blue line indicates the linear regression line between two methods and the black line indicates the identity line. The slopes of the regression lines are not significantly different from one for any pair of methods.

We performed a similar analysis for the three other iPSYCH disorders analyzed, namely ASD, Depression, and Schizophrenia. The Manhattan, Z-scores, and *χ*^2^ statistics plots can be found in **Figures S14-S19**. For depression and schizophrenia, we found no genome-wide significant hits for any method used, and the Z-scores and *χ*^2^ statistics indicate no difference in power between standard GWAS, LT-FH, and LT-FH++. For autism, we do see genome-wide significant hits, 3 for case-control GWAS and LT-FH++ and 4 for LT-FH. The SNP that is unique to LT-FH is also highly suggestive for case-control GWAS and LT-FH++. A table containing the COJO independent SNPs can be found in **Table S2 & S3** for ADHD and ASD.

## Discussion

Several large genetic datasets with linked electronic health registries (EHR) have emerged in recent years, e.g. the UK biobank data^51^, the iPSYCH data^31^, FinnGen^52^, deCODE, and many more. As more genetic data is linked to EHR, it is essential to develop statistical methods that make best use of all this information to decipher the genetics of common diseases. Here we present a new and scalable method LT-FH++ for improving power in GWAS when family history and an age-of-onset distribution is available, which is typically the case in EHR. We demonstrated the feasibility and relevance of the approach using both simulations and real data applications. Using simulated case-control outcomes with a prevalence of 5% and 10%, we observed power gains of up to 17.8% compared to LT-FH and up to 60.6% compared to using standard case-control status. We found that LT-FH++ provided the largest relative improvements when cases were ascertained (such that in-sample case-control ratio becomes larger than prevalence), and when prevalence was high.

We acknowledge that not everyone has access to the same level of detailed health records. Therefore, we would like to point out that it is not a requirement to estimate prevalence curves in the population that you are performing the analysis in. The curves can be estimated in an external population, and subsequently used to assign the personalized thresholds in the internal population provided information, such as sex, age-of-onset, and birth year, is available in the internal and external data.

We applied LT-FH++ to study mortality in UK biobank and 4 common psychiatric disorders in iPSYCH, all prevalent outcomes for which we had both family history available as well as age-of-onset distributions. This includes age, age-of-onset (for cases), cohort effects, and sex for both the genotyped individuals and family members. We also had access to public data for mortality incidence rates by age, sex and birth year for England and Wales from 1840s to the present day. We compiled similar information for the 4 psychiatric disorders using the full Danish register data (see Methods). For mortality in the UK biobank data, we found 10 independent genome-wide significant associations when applying LT-FH++, compared to 2 with LT-FH and none with the case-control status. This result further underlines the importance of including other information in GWAS. The power increase of LT-FH over case-control status highlights the importance of family history, and the power increase of LT-FH++ over LT-FH highlights the importance of accounting for age-of-onset. The most significant association was found in the APOE gene, which also harbored the only significant association in a recent survival model (frailty model) GWAS of mortality in the UK biobank data^27^. All of the identified associations were in or near well known disease-related genes and were largely concordant with the genome-wide associations found by Pilling et al.^53^ when performing a GWAS of combined mothers’ and fathers’ attained age.

We further applied LT-FH++ to the 4 common psychiatric disorders in the iPSYCH data. Combined, we found 10 independent genome-wide significant associations with LT-FH++, compared to 8 for LT-FH and case-control status. Compared to mortality, the observed power gain for the iPSYCH disorders was small, despite having access to more information per individual. The discrepancy in performance when applied to the mortality in the UK biobank and 4 common psychiatric disorders may have several reasons. First, case-control, LT-FH and LT-FH++ performed similarly for each of the 4 common psychiatric disorders, and in the simulations, we saw a relative power increase when cases were ascertained through downsampling of controls, however, due to the lower overall sample size, the absolute power to detect causal SNPs also decreased significantly with sample size. We suspect a similar situation might be happening in the iPSYCH data. Second, since simulations showed the power improvement was larger when prevalence was higher and cases were ascertained, the difference may be explained by the large lifetime prevalence difference between death and the psychiatric disorders. Third, it is possible that the multivariate liability threshold model (underlying LT-FH and LT-FH++) may better fit mortality than the psychiatric disorders. More specifically, the model makes several key assumptions. First, both LT-FH and LT-FH++ assumes that the heritability is known and that there is no environmental covariance between family members. In practice, one can often estimate the heritability in the sample or rely on published estimates. Second, it assumes that the population disease prevalence is known, and (if relevant) provided for subgroups defined by age, birth year, and sex. However, simulations using LT-FH and LT-FH++ indicate that it is relatively robust to misspecification of these parameters^30^. Third, the model assumes that the genetic architecture of the disease or trait in question does not vary by age of diagnosis, birth year, or differ between sexes. Some research suggests that this assumption is reasonable for many outcomes, including the four psychiatric disorders analysed here^54,55^, but these will generally not hold in practice. We note that case-control GWAS also assume this unless the analysis is stratified by these subgroups. Fourth, LT-FH++ assumes that the threshold always decreases with age. The intuition behind this is that the disease prevalence is the cumulative incidence, which by definition always increases with age, and the threshold is the upper quantile of the inverse standard normal at the age specific prevalence. An individual then only becomes a case if its liability becomes larger than the prevalence threshold, as it decreases with time. A consequence of this assumption is that early-onset cases generally have higher disease liabilities than late-onset cases, which is also the expectation in survival model analysis if the hazard rate is (positively) correlated with the genetic risk. The correlation between genetic risk and earlier age-of-onset has been observed for several common diseases, e.g. Alzheimer’s disease^56^, coronary artery disease and prostate cancer ^57^. However, if the age-of-onset for a given disease is not heritable, or if the genetic correlation between the age-of-onset and disease outcome is weak, then we do not expect LT-FH++ to improve statistical power for identifying genetic variants. Indeed this might be one possible explanation for why we do not observe improvements in power when applying LT-FH++ to iPSYCH data, although we note that polygenic risk scores have been found to contribute to hazard rates for psychiatric disorders in the iPSYCH data^58,59^.

Conceptually, LT-FH++ combines two methods into one to improve power in genetic analyses, namely LT-FH, which is based on the liability threshold model and incorporates family history, and survival analysis, which can account for age and changes in prevalence over time and is routinely used to model time-to-event data. With family history and age-of-onset information available, we believe LT-FH++ will be an attractive method for improving power in many different genetic analyses, including GWAS, heritability analyses and for polygenic risk scores^60–62^. As more genetic datasets with linked health records and family information become available, e.g. in large national biobank projects, we expect the value of statistical methods that can efficiently distill family history and individual health information into biological insight will only increase.

## Acknowledgements

We would like to thank Margaux Hujoel for useful discussions and allowing us to use the LT-FH++ name. We would like to thank Mark Daly for useful advice and helpful comments. F.P. and B.J.V. were supported by the Danish National Research Foundation (Niels Bohr Professorship to Prof. John McGrath). We also acknowledge the Lundbeck Foundation Initiative for Integrative Psychiatric Research, iPSYCH (R102-A9118, R155-2014-1724 and R248-2017-2003). B.J.V. was also supported by a Lundbeck Foundation Fellowship (R335-2019-2339). High-performance computer capacity for handling and statistical analysis of iPSYCH data on the GenomeDK HPC facility was provided by the Center for Genomics and Personalized Medicine and the Centre for Integrative Sequencing, iSEQ, Aarhus University, Denmark (grant to A.D.B.).

## Code Availability

The code used for LT-FH++ has been implemented into an R package, and it is available at https://github.com/EmilMiP/LTFHPlus. We have also reimplemented LT-FH in the package, where we utilize the Gibbs sampler to efficiently estimate the genetic liabilities, keeping the same input format as the original implementation.

## Methods

### Model

The underlying model is identical to the one used in LT-FH^30^, as a result the model will only briefly be presented here, and the main differences will be elaborated on. Under the liability threshold model each individual has a liability, *l*, which follows the standard normal distribution. An individual will be considered a case, *z* = 1, when their liability is above a given threshold, i.e. *l* ≥ *T*, and a control, *z* = 0, if the liability is below the threshold, *l* < *T*. The threshold, *T*, is determined from the prevalence of the dichotomous disorder, such that *P*(*l* ≥ *T*) = *K*, where *K* denotes the prevalence in the population.

LT-FH builds on this idea, and for a single individual, the liability is assumed to be further decomposed into a genetic and environmental component, *l* = *l_g_* + *l_e_*. Both *l_e_* and *l_g_* are normally distributed and independent. We have:

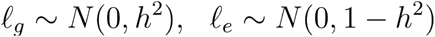

Here *h*^2^ is the heritability on the liability scale. The LT-FH setup extends this idea to include parents and siblings. It considers a multivariate normal distribution given by:

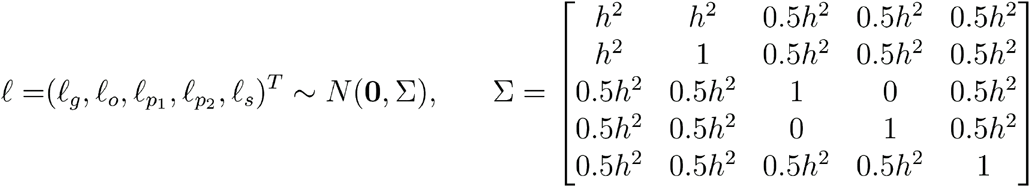

Here *ℓ_o_* denotes the full liability for the individual (denoted *ℓ* for a single individual above), and *ℓ_g_* denotes the genetic component of this liability. *ℓ*_*p*1_ and *ℓ*_*p*2_ denotes the *full* liability of each parent, while *ℓ*_S_ denotes those of the sibling. The example above includes one sibling only, but in theory any number of siblings could be included in the model. We are interested in estimating the posterior mean genetic liability for each individual conditional on family information:

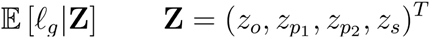

Here **Z** denotes the vector of status for the family, consequently a restriction is placed on each individual’s full liability. In the case of everyone having the disorder, we would consider the space {*ℓ* ∈ ℝ|*ℓ_i_* ≥ *T_i_* for all *i*}, where denotes a family member, and *T_i_* denotes the family member’s threshold and *ℓ_i_* denotes their full liability. In LT-FH, the thresholds are the same for all children (the offspring and any siblings), and another threshold is used for all parents.

The choice of thresholds is where LT-FH++ starts to differentiate itself from LT-FH. In short, the liability thresholds are personalized, such that every individual, sibling or parent has a potentially unique threshold which is determined by their age, birth year, and sex. Furthermore, we adapt an age-dependent liability threshold model, where the threshold is dynamic, in the sense that it decreases as a population grows older. This idea is illustrated in figure 1A, where the threshold decreases as time progresses for a population, with marks for ages 15, 25, 35, 50, and 80. This model assumes that the threshold decreases continuously as time progresses, and these marks can be seen as snapshots in time, where an individual who was diagnosed at one of the marks had an assumed (fixed) liability equal to said mark. This age-dependent liability threshold model allows us to be very precise with the liability for cases when an accurate age-of-onset is available. If an accurate estimate of age-of-onset is not available, then the threshold can still be personalized based on other available information, with the modification that we do not fix the full liability, but integrate over all liabilities above the personalized threshold. Interestingly, the age-dependent liability threshold model can be thought of as a survival analysis (see below).

Another point where LT-FH++ differs from LT-FH is in how siblings are included. LT-FH includes the siblings by specifying the number of siblings and assigns a single case-control status to the siblings with the condition that at least one sibling has the disorder. However, a more fine-grained inclusion of the siblings, where each sibling is added individually is not available. LT-FH++ expects each individual and their family members to be added separately, such that information on each individual can be accounted for.

### Relationship with survival analysis

In survival analysis GWAS the risk for becoming a case in a time-interval depends on the covariates in the model. This is reflected by a hazard rate *λ*(*t*|*x*), which describes the event rate. In our context it would refer to the rate for becoming a case. This rate depends on both time *t*, and covariates of the model *x*, e.g. genotypes. The hazard rate (also referred to as the intensity) can be approximated by 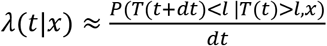, where *dt* is a small change in time^63^, *T*(*t*) is the threshold for being a case at time *t*, and *l* is the full liability of an individual. This means that the hazard rate is proportional to the probability an event occurs within a time-interval (*t*, *t* + *dt*), given that no event had occurred earlier. For different types of survival analyses we can estimate this probability using the hazard rate, e.g. for a Cox proportional hazards model where we aim to estimate the effect of a genotype *x* on the hazard rate, it becomes: *P*(*T*(*t* + *dt*) < *l* |*T*(*t*) > *l*, *x*) = *dt λ*(*t*|*x*) = *dt λ*_0_(*t*)*exp*(*βx*). To keep notation simpler, we will denote the genetic liability of individual *i* as *g_i_* instead of *l_gi_*, and if we further assume that the genetic component for an individual of a case-control outcome contributes to the hazard rate such that *λ*(*t*|*g_i_*) = *λ*_0_(*t*)*exp*(*g_i_*) = *λ*_0_(*t*)*exp*(*β x_i_*), where *x_i_* denotes the genotype of the i’th individual, and *β* their true effects (in the Cox-regression model). Conceptually, this means that individuals with higher than average genetic risk, i.e. *g_i_* > 0, will be at higher risk to be come cases throughout their lives, irrespective of age. These high-risk individuals will on average also have earlier age-of-onset.

To understand how this model relates to the proposed age-dependent liability threshold model, we can derive the same probability to approximate the corresponding hazard rate. Under the LT-FH++ model, the probability for an individual *i* to be diagnosed (become a case) within a time-interval *dt* can be written as *P*(*T*(*t* + *dt*) ≤ *l_i_*|*T*(*t*) > *l_i_*, *g_i_*), where *t* again denotes the age of the individual, and *T*(*t*) now denotes the age-dependent liability threshold. We note that *T*(*t*) is a monotonic decreasing function as the prevalence of a case-status (i.e. cumulative lifetime incidence proportion) always increases with age (conditional on birth year and sex). Furthermore, *l*_*i*_ denotes the full liability of the individual and *g_i_* the genetic component of that liability (which is generally on a different scale than a genetic component in Cox regression). The liability threshold model assumes that the liability of an individual consists of a genetic and environmental components, i.e. *l_i_* = *g_i_* + *e_i_*. It also assumes that these are in dependent, follow a Gaussi an distribution, and have variance *h*^2^ and 1 − *h*^2^ respectively. Hence using these we can expand the probability of being diagnosed within a time-interval further as follows

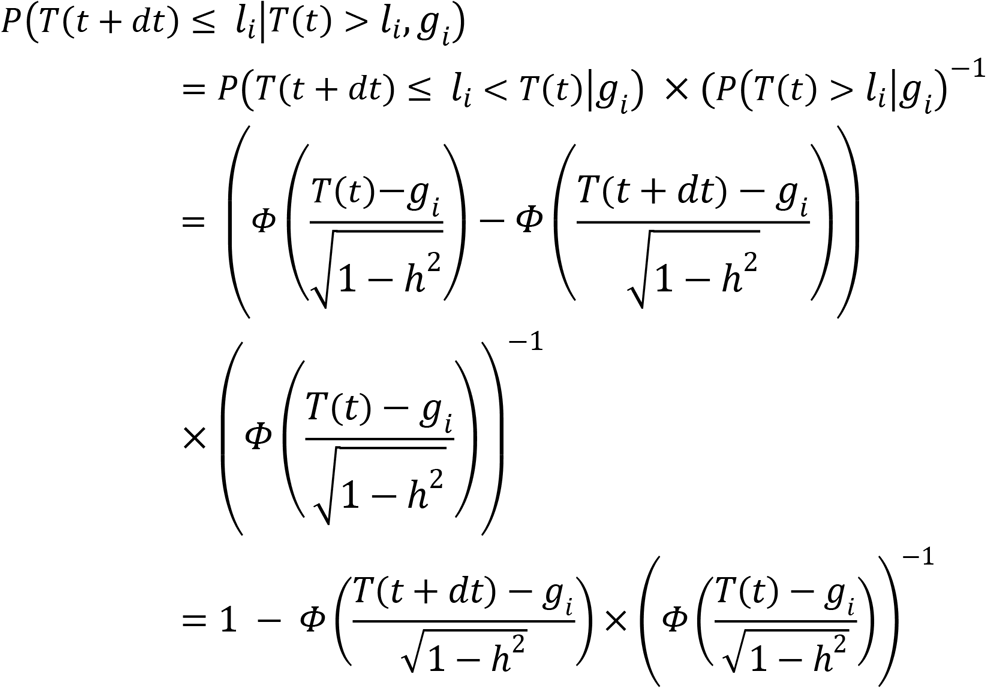

Plotting this function for different thresholds and genetic liability values shows that the probability for being diagnosed within the time-interval, and thus the hazard rate, increases linearly as a function of the genetic liability when *g_i_* is near *T*(*t*) or larger. We compare this probability with the corresponding Cox regression probability assuming a base incidence rate of *λ*_0_(*t*) = *α*, where *α* is determined by the prevalence. These two probabilities, which are proportional to the hazard rate, are plotted as a function of *g_i_* in **Figure S20**, illustrating how the hazard rates of the two models depend on *g_i_*. We note that the two models share the properties that individuals with higher than average genetic risk will, on average, be more likely to become cases within any time-interval, and have earlier age-of-onset.

It may seem counterintuitive that a arguably deterministic model such as the age-dependent liability threshold model, where the liability is constant throughout life, can be recast as a survival analysis model. The reason for this is that although the outcome of the age-dependent liability threshold model is always known given the liability, one never observes this liability. Hence, the environmental term, which can be thought of as capturing various environmental effects as well as chance events and other non-genetic effects, leads to a non-deterministic survival analysis model.

### Sampling Strategy

If we consider an individual with disease status available for both parents, but no siblings, then we have a total of 6 unique ways to configure the status vector, **Z**, when disregarding other information, since the scenario where a single parent is a case can happen in two ways. LT-FH estimates the posterior mean genetic liability for each of these configurations by sampling a large number of observations from the multivariate normal distribution described above. The observations are then grouped into these 6 unique configurations, and the genetic liabilities are estimated by averaging genetic liabilities within each configuration. This strategy works well when there are a limited number of configurations, but becomes infeasible when the number of configurations becomes too large.

LT-FH++ cannot efficiently use the same sampling strategy, since the personalized thresholds increases the number of potential configurations such that the strategy becomes intractable. Instead LT-FH++ considers each family as a unique configuration, since it uses individualized thresholds. To derive the posterior means efficiently we use a Gibbs sampler to sample from a truncated multivariate normal distribution^64^. The truncation points in the truncated multivariate normal distribution are the personalized thresholds. Sampling for all individuals is fast, requires far fewer observations, and can be easily parallelized across individuals as each family is independent from each other.

### Prevalence Information

The age-dependent prevalence of ADHD, autism, depression and schizophrenia was obtained through Danish national population-based registers. For these estimates, we included all 9,251,071 persons living in Denmark at some point between January 1, 1969 and December 31, 2016. Each individual in the study was followed from birth, immigration to Denmark, or January 1, 1969 (whichever happened last) until death, emigration from Denmark, or December 31, 2016 (whichever happened first). All dates were obtained from the Danish Civil Registration System^65^, which has mantained information on all residents since 1968, including sex, date of birth, continuously updated information on vital status, and a unique personal identification number that can be used to link information from various national registers. Information on mental disorders was obtained from the Danish Psychiatric Central Research Register^66^, which contains data on all admissions to psychiatric inpatient facilities since 1969 and visits to outpatient psychiatric departments and emergency departments since 1995. The diagnostic system used was the Danish modification of the *International Classification of Diseases, Eighth Revision (ICD-8)* from 1969 to 1993, and *Tenth Revision (ICD-10)* from 1994 onwards. The specific disorders were identified using the following ICD-8 and ICD-10 codes: ADHD (308.01 and F90.0), autism (299.00, 299.01, 299.02, 299.03 and F84.0, F81.4, F84.5, F84.8, F84.9), depression (296.09, 296.29, 298.09, 300.49 and F32, F33), and schizophrenia (295.x9 excluding 295.79 and F20). For each individual in the study, the date of onset for each disorder was defined as the date of first contact with the psychiatric care system (inpatient, outpatient, or emergency visit). All analyses were done separately for each sex and for each birth year. The cumulative incidence function for each disorder was estimated with the Aalen-Johansen approach considering death and emigration as competing events^67^. The cumulative incidence over age is interpreted as the proportion of persons diagnosed with the specific disorder before a certain age.

### Personalized Thresholds

With the cumulative incidence rate tables we are able to assign personalized thresholds to everyone with sufficient information available. Examples of cumulative incidence rate curves can be seen in **Figures S9 - S12**. Under the liability threshold model, sex, birth year and age for controls or age-of-onset for cases can uniquely determine the threshold for an individual. Based on this information a proportion is assigned to them, which is transformed to an individual’s threshold through the inverse normal cumulative distribution function.

For controls, it has allowed us to tailor the threshold in the liability threshold model to each individual, similar to what is seen in **Figure 1A**, where the threshold is decreasing as an individual is getting older. In short, the older a control is, the larger a proportion of the possible liabilities in the liability threshold model can be excluded as no longer attainable. For cases, the tailored threshold means we are able to very accurately estimate what a person’s *full* liability is for a given disorder under the liability threshold model. Since the full liability can be accurately estimated for a case by the assigned threshold, we will fix the full liability of a case to be the threshold in the model.

### Simulation details

For the simulations we simulated 100.000 unrelated individuals each with 100.000 independent single-nucleotide polymorphisms (SNPs). We simulated two parents, and between zero and two siblings. The parents’ genotypes were drawn from a binomial distribution with probability parameters equal to the allele frequency (AF) of the corresponding variant. The variant AF was drawn from a uniform distribution on the interval (0.01, 0.49). The parents’ genotypes were either 0, 1, or 2; we defined the child’s genotypes as the average between the genotypes of both parents, rounding values of 0.5 or 1.5 up or down with equal probability. Allele effect sizes were drawn from *N*(0, *h*^2^/*C*), with *C* being the number of causal SNPs and *h*^2^ denoting the heritability. Case-control status was assigned using the liability threshold model.

The default simulation setup consisted of causal SNPs assigned to positions at random, two different prevalences, 5% and 10%, *C* set to 1000, and a sex-specific prevalence of 8% for men and 2% for women. When the prevalence was 10%, these sex-specifc prevalences were doubled. To generate the age-of-onset, we assumed that the cumulative incidence curve followed a logistic function, because it resembles real-world cumulative incidence rates for some traits, see **Figure S7 & S10-13**. The logistic function is given by:

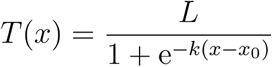

where *L* denotes the maximal attainable value, *k* is the growth rate, and *x*_0_ denotes the age at which *K* is *L*/2, which is the midpoint of the curve, i.e. median age-of-onset. These parameters resulted in an age-of-onset that was largely normally distributed around the median age, *x*_0_. The cumulative incidence rate curve allows us to obtain the expected prevalence at each age, which we can then translate into a threshold in the liability threshold model, i.e. an earlier diagnosis indicates higher liability for the trait. We set *L* as the lifetime prevalence (5% and 10%). We also set *k* to 1/8 and *x*_0_ to 60 such that 90% of cases have an age-of-onset between 36.5 and 83.5.

A family consisted of one offspring, two parents, and zero to two siblings. The age of the cases was set to the age-of-onset. The age-of-onset was assigned by taking the inverse of the logistic function on the full liability’s quantile under the standard normal distribution. Individuals with an age lower than their age-of-onset would normally be considered controls, since they had not yet had the time to develop the disorder. However, setting high liability individuals to controls due to age-of-onset being later than age was decided against to properly fix the number of cases to the prevalence in the simulated data. For controls, the offspring’s age was uniformly distributed between 10 and 60. The parents’ age was set to the age of the child plus a uniform draw between 20 and 35, allowing for up to 95 year olds. The threshold was assigned with the logistic function with the age and sex as inputs. For simplicity, birth year was not modelled.

When considering sex and the sex-specific prevalences, we assigned each individual a male or female sex with equal probaility. We assumed males were four times as likely to be cases than females. For the two overall prevalences (5% and 10%) this corresponded to 8% and 16% prevalence among males (liability thresholds *T*_*male*_ = 1.41 and *T*_*male*_ = .99) and 2% and 4% prevalence among females (liability thresholds *T*_*female*_ = 2.05 and *T*_*female*_ = 1.75). Finally, we simulated sample ascertainment by downsampling controls such that cases and controls had equal proportions (50% each). For 5% prevalence this resulted in a sample size of 10,000 and 20,000 individuals when using a prevalence of 10%.

### GWAS in UK biobank

We restricted individuals to the White British group (Field 22006) and to the individuals used for computing the principal components (PCs) in the UK Biobank (Field 22020). These individuals are unrelated and have passed some quality control (see section S3 of ^51^). This resulted in 337,475 individuals. **Table 1** shows a breakdown of how many people are cases and controls for the genotyped individuals and parents. We used the genotyped SNPs for the UK biobank participants as model SNPs in BOLT-LMM^37^, after removing SNPs with maf < 0.01, missing call rate > 0.01, and Hardy-Weinberg equilibrium p-value < 1 × 10 ^−50^, which left us with a total of 504,138 SNPs. When performing the GWAS, we used the imputed SNPs in bgen files and removed SNPs with a maf < 0.005 or info score < 0.6, which resulted in 11,335,564 SNPs. We used BOLT-LMM v2.3.2 with age, sex, and the first 16 PCs as covariates. The three mortality outcomes used in the UK biobank were case-control status, LT-FH, and LT-FH++. We considered the binary death outcome as the case-control phenotype, with LT-FH and LT-FH++ further utilising the mortality status of both parents, but no siblings.

**Table 1:** Breakdown of the number of cases and controls for mortality for the UK biobank participants (here Children) and their parents. The case-control GWAS only used the Children column as input, while LT-FH and LT-FH++ used all columns.

| Mortality |  |  |  |  |  |
| --- | --- | --- | --- | --- | --- |
| Children |  | Father |  | Mother |  |
| Case | Control | Case | Control | Case | Control |
| 13,819 | 323,656 | 258,932 | 75,545 | 199,856 | 130,757 |

LT-FH++ and LT-FH require prevalence information, which was acquired from the Office for National Statistics (ONS) https://www.ons.gov.uk/. Mortality rates for England and Wales were available from 1841 to the present day. The same information was available for all of the United Kingdom (UK), but only from the 1950’s onwards. Since England is the most populous country in the UK, we believe these mortality rate estimates are a good proxy for all of the UK. From the mortality rates provided by ONS, we calculated the cumulative incidence curves for death, for each birth year from 1841 onwards and for both sexes. We used this information to calculate the personalized thresholds in LT-FH++, accounting for birth year, sex, and current age or age of death. Note that, in LT-FH, it is not possible to adjust for sex, age, or cohort effects at the individual level, but two different threshold can be specified, one for all parents and one for all children. Therefore, we assumed the same age for all children and the same age for all parents when running LT-FH. We used the last recorded death as the endpoint, which happened in 2018, and assumed all children were 55 years old, and parents were 85 years old. This translated into an assumed birth year of 1963 and 1933, respectively. Based on these birth years, we found the prevalence of death for these birth years and ages in the survival curve, and averaged the sex-specific prevalences. For LT-FH, we also considered thresholds based on prevalence estimated in the UK biobank participants and their parents, however we did not see any significantly different results, when comparing to the population based prevalence estimates (results not shown). A heritability of 20% was used for LT-FH and LT-FH++.

### GWAS in iPSYCH

The iPSYCH cohort has recently received a second wave of genotyped individuals, increasing the number of genotyped individuals from ~80k to ~143k^68^. The two iPSYCH waves have been imputed separately with the Ricopili pipeline^69^. After combining the two waves and removing any SNP with missingness > 0.1 or MAF < 0.01, we have a total of 4,706,774 SNPs. When performing a GWAS, we restrict the analysis to individuals classified as controls in the iPSYCH design and individuals diagnosed with the analysed phenotype, even when using LT-FH or LT-FH++. We filtered for relatedness with a 0.0884 KING-relatedness cutoff and restricted the analysis to a genetically homogeneous group of individuals by calculating a Mahalanobis distance based on the first 16 PCs and keeping individuals within a log-distance of 4.5^70^. For a breakdown of the number of individuals included in each GWAS and the number of cases and controls see **Table 2**. We used BOLT-LMM^37^ v2.3.2 to perform the GWAS with sex, age, wave, and the first 20 PCs as covariates. LT-FH and LT-FH++ require an estimate for the heritability, and for ADHD we used 75%^71^, autism was 83%^72^, depression was 37%^73^, and schizophrenia was 75%^73,74^. See Prevalence Information for details on how the cumulative incidence curves were derived.

**Table 2:** Breakdown of how many cases and controls each GWAS was performed with. For case-control outcome only the Children column was used. For LT-FH and LT-FH++ all columns were used. LT-FH only included a binary variable for sibling status, for ASD this meant 1,427 satisfied the “at least one sibling is a case” condition of LT-FH. Some individuals had no siblings and thus no sibling status.

|  | Children |  | Father |  | Mother |  | Sibling Status |  |  |  |
| --- | --- | --- | --- | --- | --- | --- | --- | --- | --- | --- |
|  | Case | Control | Case | Control | Case | Control | 0 | 1 | 2 | 3 |
| ADHD | 21,255 | 36,584 | 498 | 57,001 | 751 | 57,088 | 43,558 | 1,777 | 78 | < 5 |
| ASD | 18,076 | 36,781 | 84 | 54,438 | 76 | 54,781 | 42,585 | 1,359 | 62 | 6 |
| Depression | 27,266 | 38,882 | 2,632 | 63,164 | 4,336 | 52,821 | 50,449 | 2,281 | 86 | 5 |
| Schizophrenia | 5,749 | 36,961 | 429 | 42,051 | 576 | 34,871 | 34,358 | 494 | 16 | < 5 |

When assessing power between outcomes, we considered SNPs that are in the iPSYCH cohort and have been found to be significantly associated with the psychiatric disorder being analysed in the largest publicly available meta-analysed GWAS^8–10,75^. PLINK is used to perform linkage disequilibrium (LD) clumping on the external summary statistics. We used PLINK’s default parameters, except for the significance thresholds. PLINK’s p-value threshold used were 5 × 10^−6^ for both the index SNP and the clumped SNPs. The default window size of 250kb and the LD threshold of 0.5 was used.

