## Supplemental Figures and tables for "Accounting for age-of-onset and family history improves power in genome-wide association studies"

### Supplementary Information

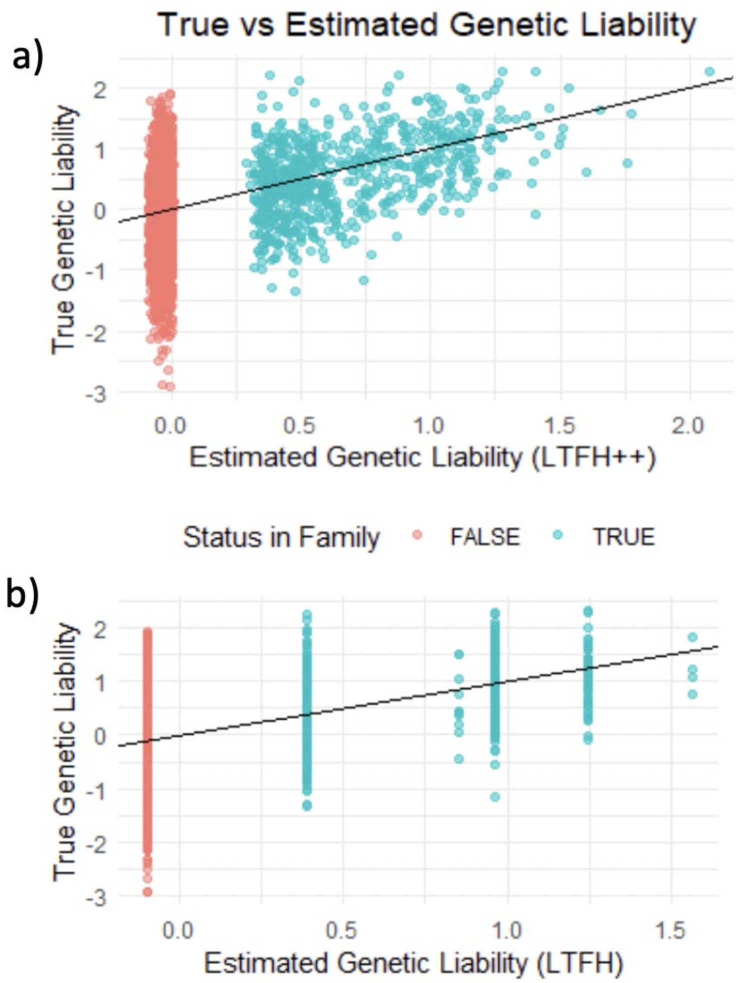


Figure S1: Simulated genetic liabilities assuming two parents and 0 siblings, a heritability of 50% and a prevalence of 10% (see Methods for details). In b) we see that LT-FH estimates for the genetic liabilities fall into specific groups, dependending on the case status of the individual and family members. In a) LT-FH++ takes age into account to obtain a more refined prediction of the genetic liability.

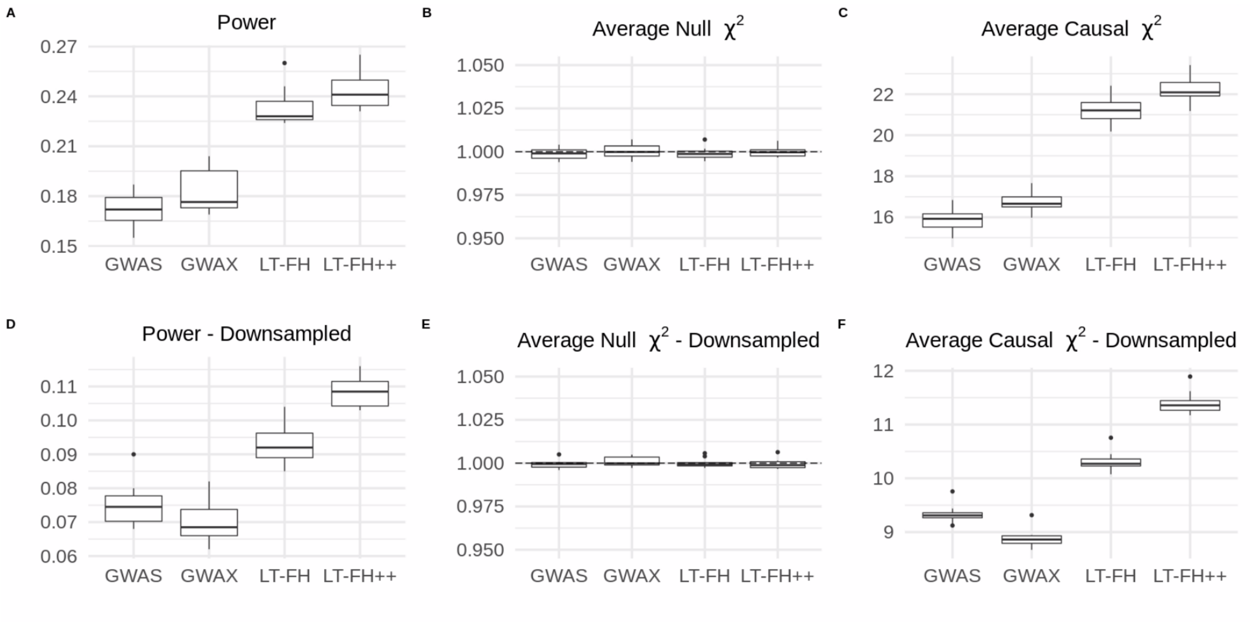


Figure S2: Simulation results under the default simulation parameters and a prevalence of 10%.

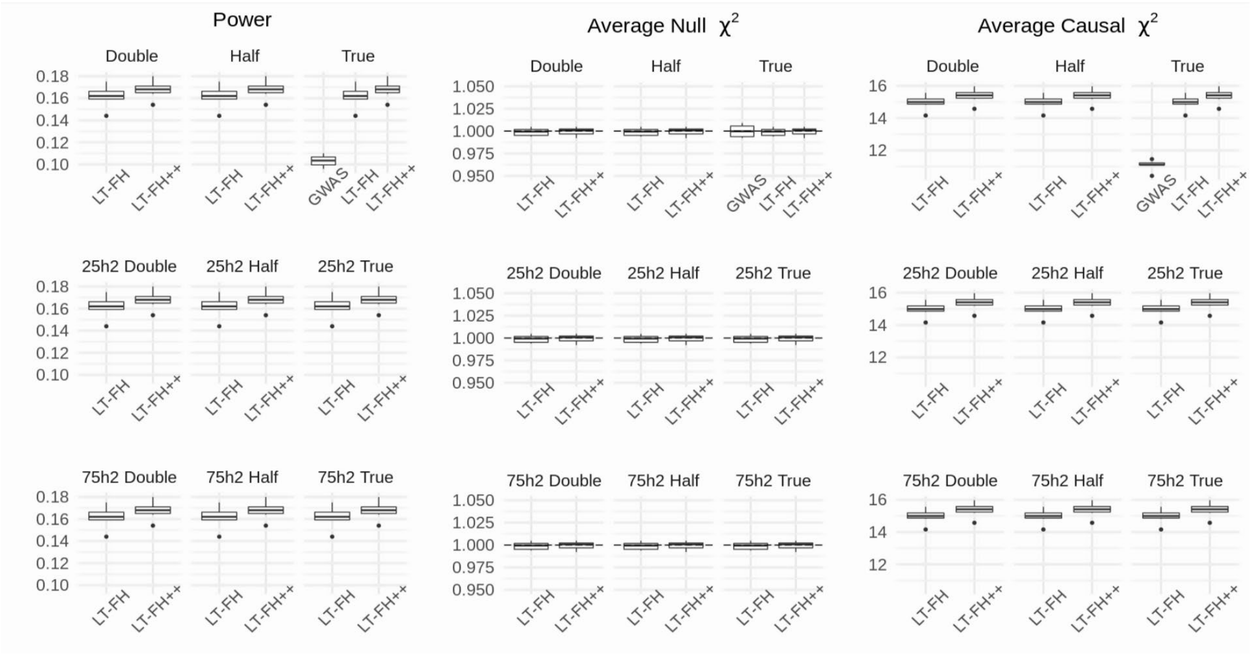


Figure S3: Simulation results with misspecified parameters and a prevalence of 5%. “Half” and “Double” refers to the misspecified prevalence, where “Half” means half of the true prevalence was used, and “Double” means double of the true prevalence was used. For reference, we added “True”, which is the true prevalence.

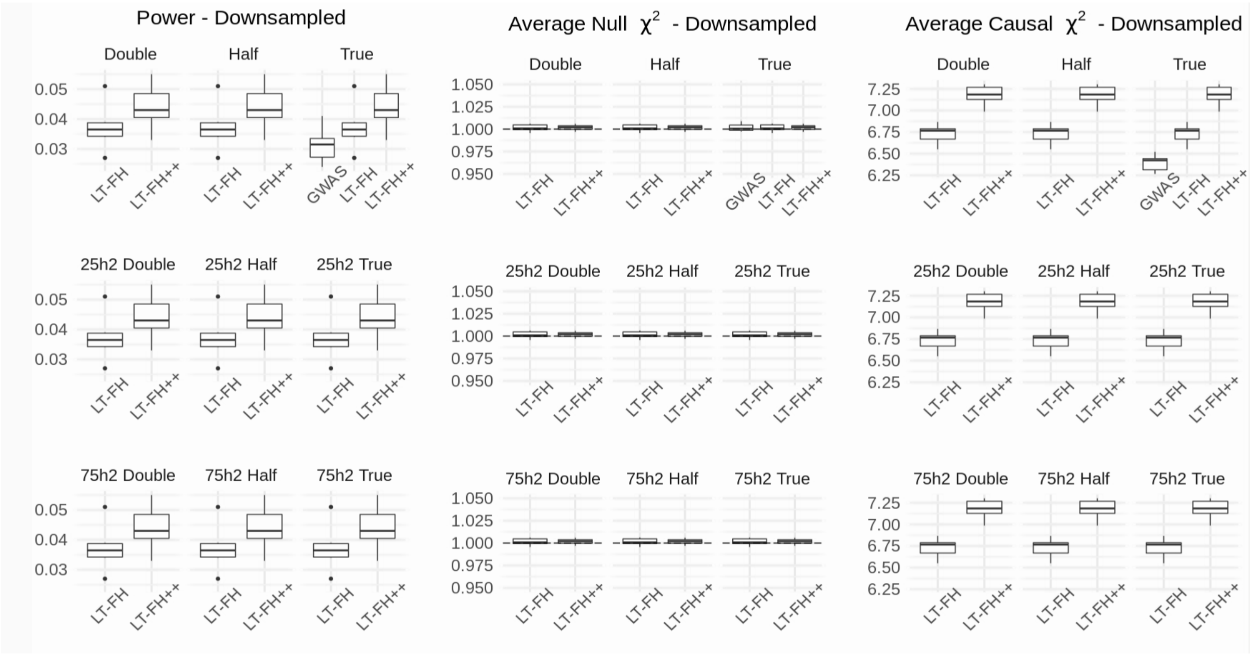

Figure S4: Simulation results with misspecified parameters, a prevalence of 5%, and downsampling of controls. “Half” and “Double” refers to the misspecified prevalence, and “Half” means half of the true prevalence was used, and “Double” means double of the true prevalence was used. For reference, we added “True”, which is the true prevalence.

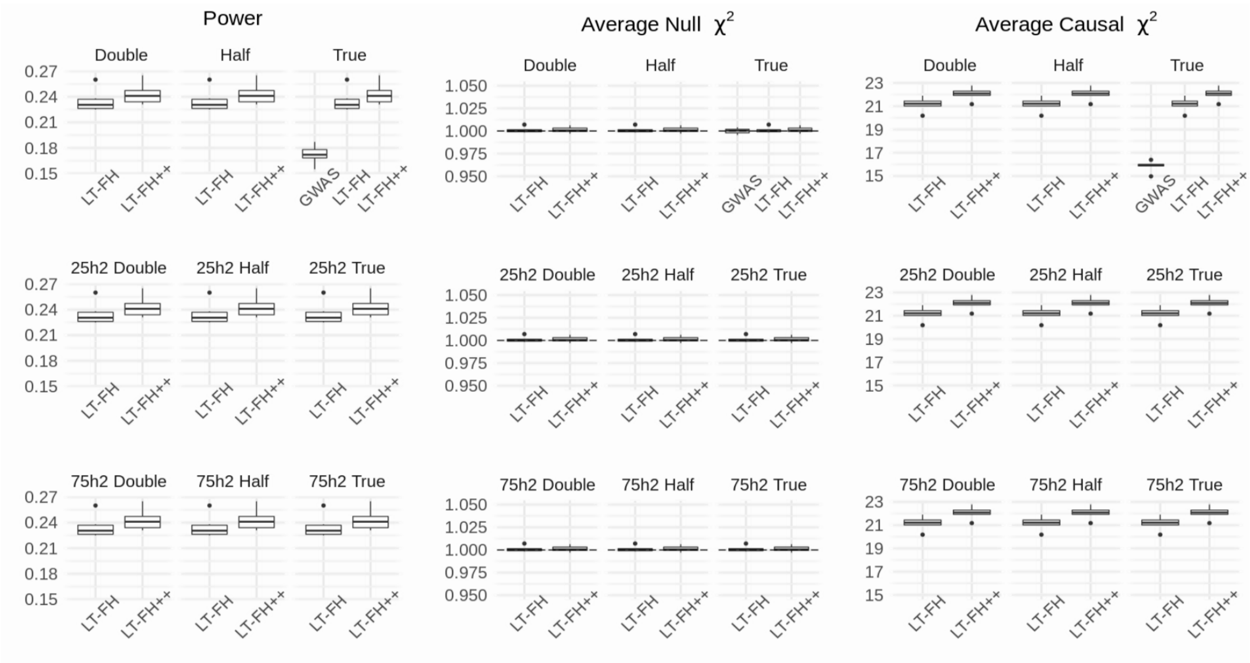


Figure S5: Simulation results with misspecified parameters and a prevalence of 10%. “Half” and “Double” refers to the misspecified prevalence, and “Half” means half of the true prevalence was used, and “Double” means double of the true prevalence was used. For reference, we added “True”, which is the true prevalence.

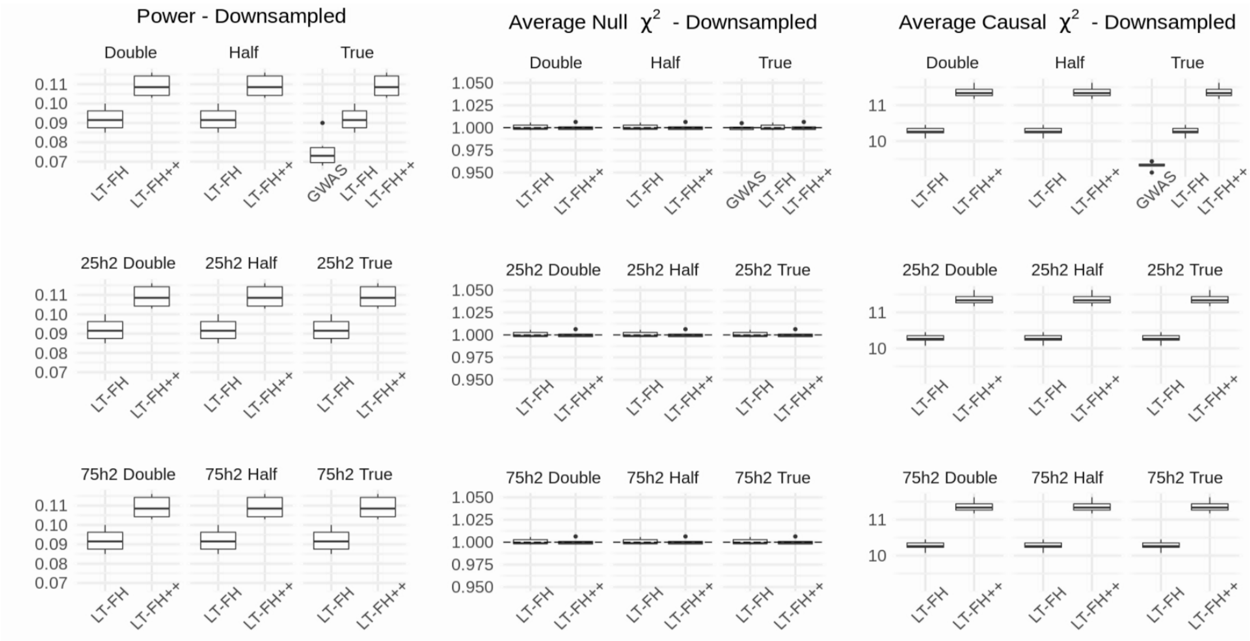


Figure S6: Simulation results with misspecified parameters, a prevalence of 10%, and downsampling of controls. “Half” and “Double” refers to the misspecified prevalence, and “Half” means half of the true prevalence was used, and “Double” means double of the true prevalence was used. For reference, we added “True”, which is the true prevalence.

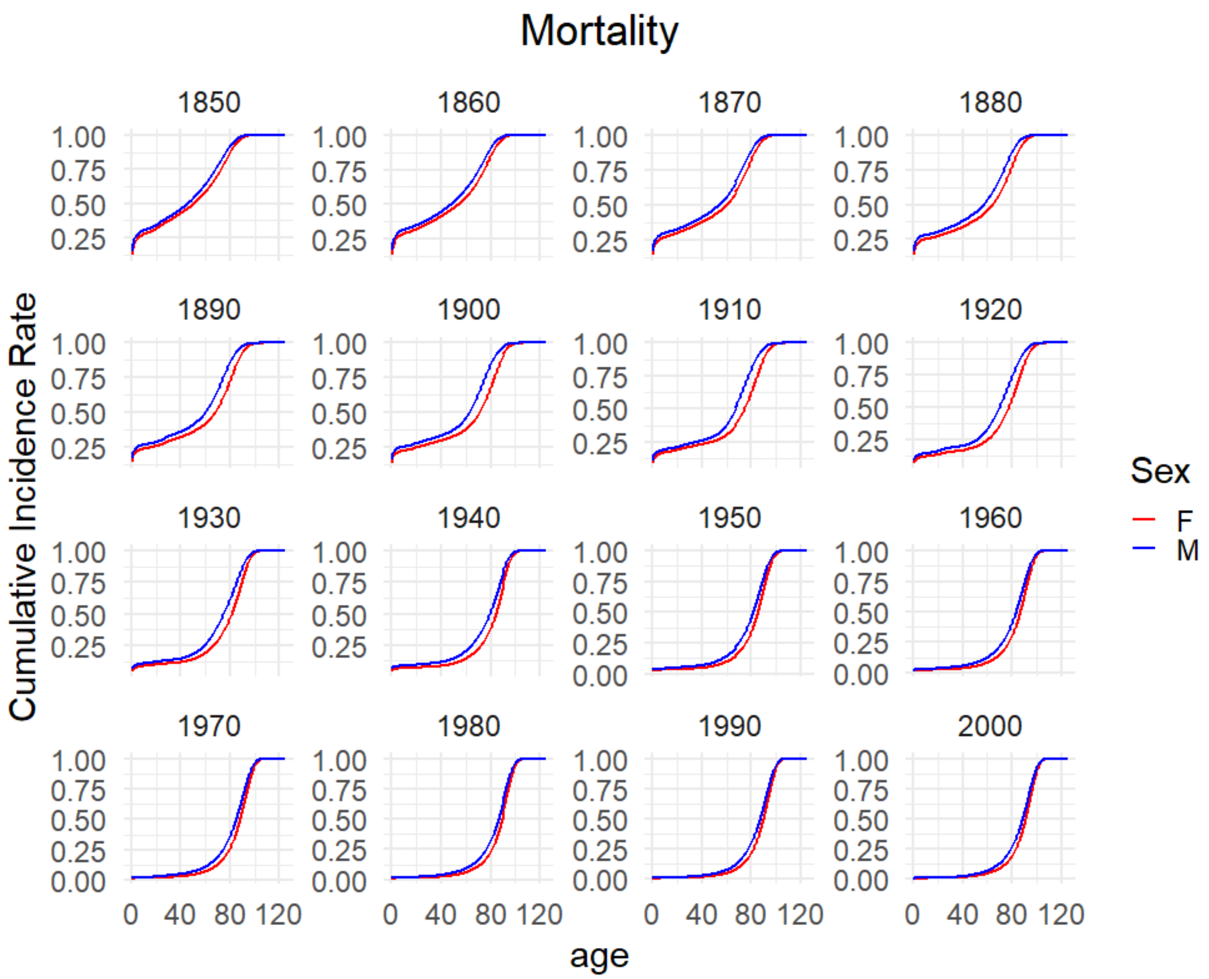


Figure S7: Plot of mortality from England and Wales, obtained from the Office for National Statistics (ONS). We plotted the cumulative mortality for each sex and from the begining of each decade from 2000 to the begining of the data. Historic mortality rates have been used upto the present, and projections for future predictions.

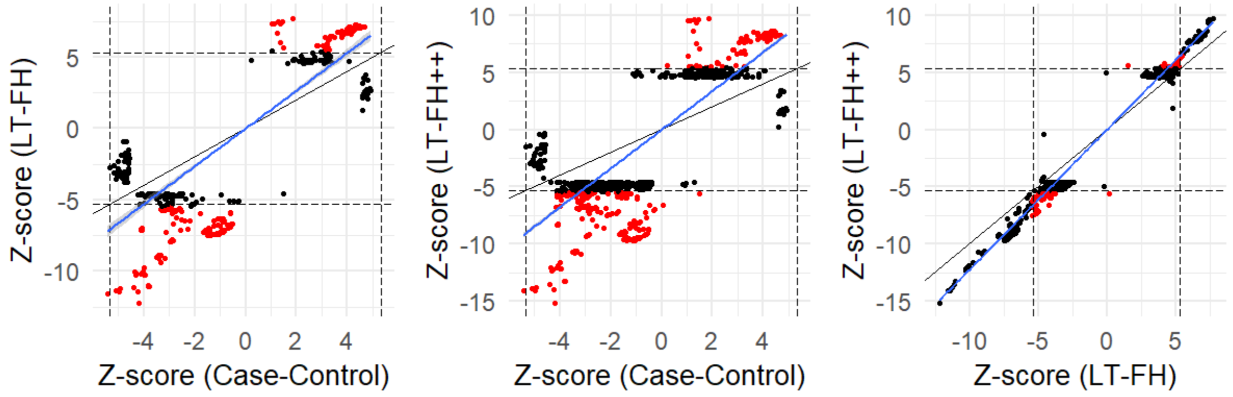


Figure S8: Z-scores for mortality in the UK biobank. The dashed line correspond to a p-value of $5 \times 10^{-8}$, and the red dots are SNPs that are genome-wide significant for only one method. The black line is the identity line and the blue line is the best fitted line. We filtered on the p-values, keeping SNPs that are below $5 \times 10^{-6}$ for at least one of the compared methods . The squared slope of the fitted line indicates the power improvement of one method over another

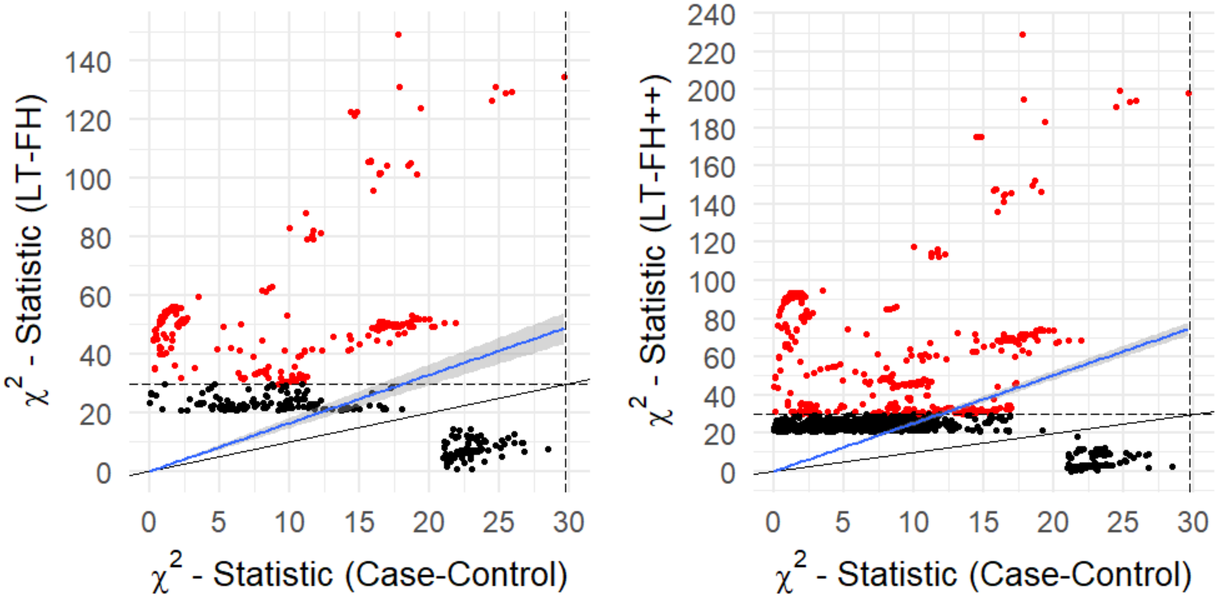


Figure S9: The $\chi^2$ statistics for mortality between case-control status and LT-FH and LT-FH++ can be seen above. The red dots are SNPs identified as genome-wide significantly associated by only one of the methods. The black dots are suggestive associations identified by either method. The black line indicates the identity line and the blue line is the best fitted line using linear regression. The black dashed lines correspond to the threshold for genome-wide significance.

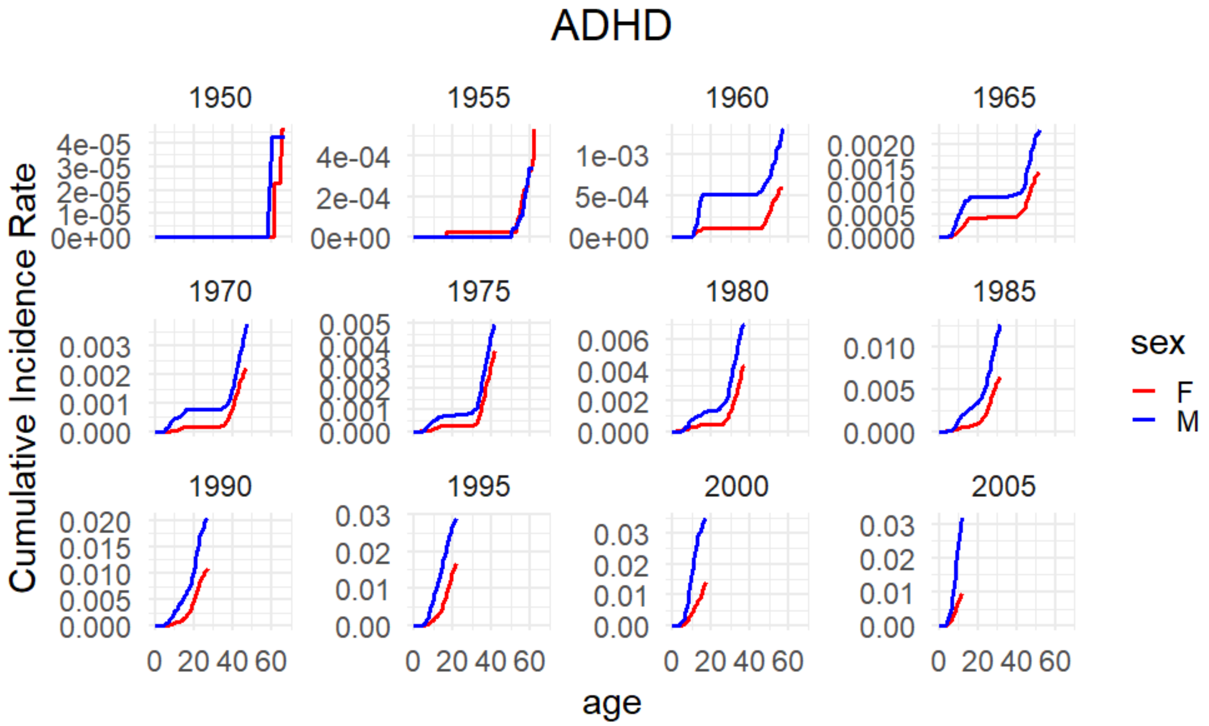

Figure S10: Plot of the cumulative incidence rate for Attention Deficit Hyperactivity Disorder grouped by birth year in the Danish registers. The red line corresponds to females and the blue corresponds to males.

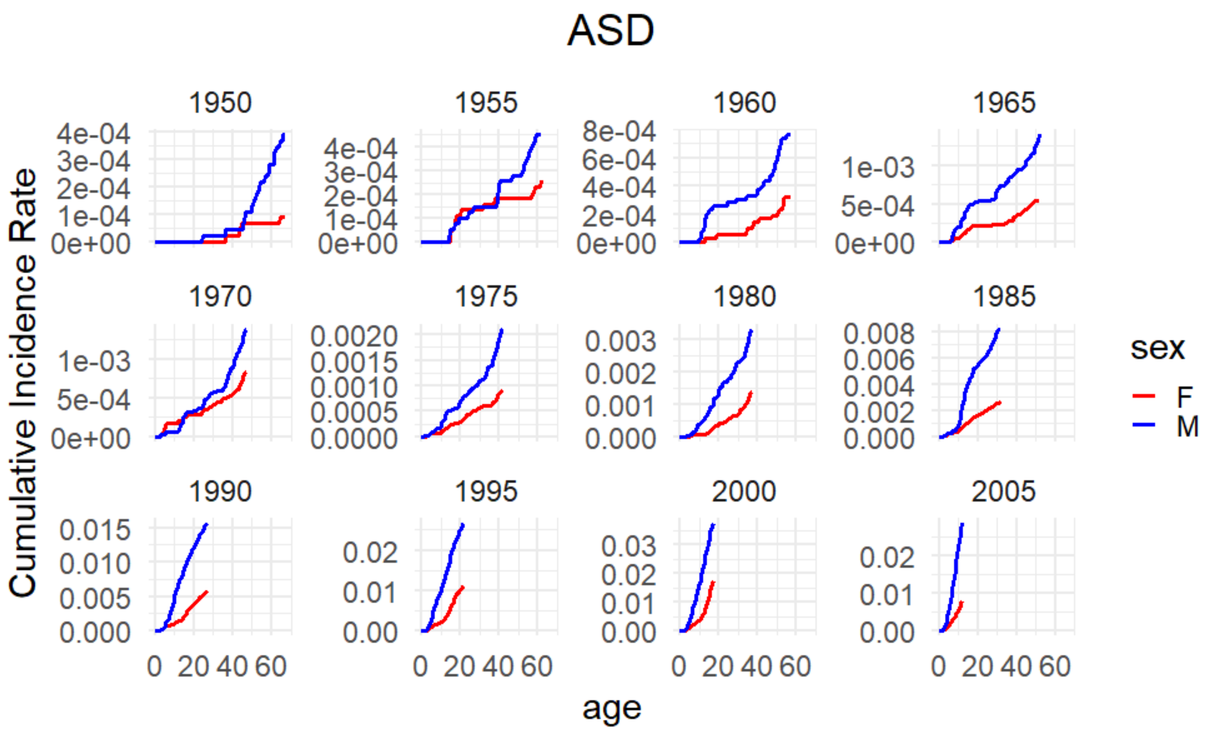


Figure S11: Plot of the cumulative incidence rate for autism spectrum disorder grouped by birth year in the Danish registers. The red line corresponds to females and the blue corresponds to males.

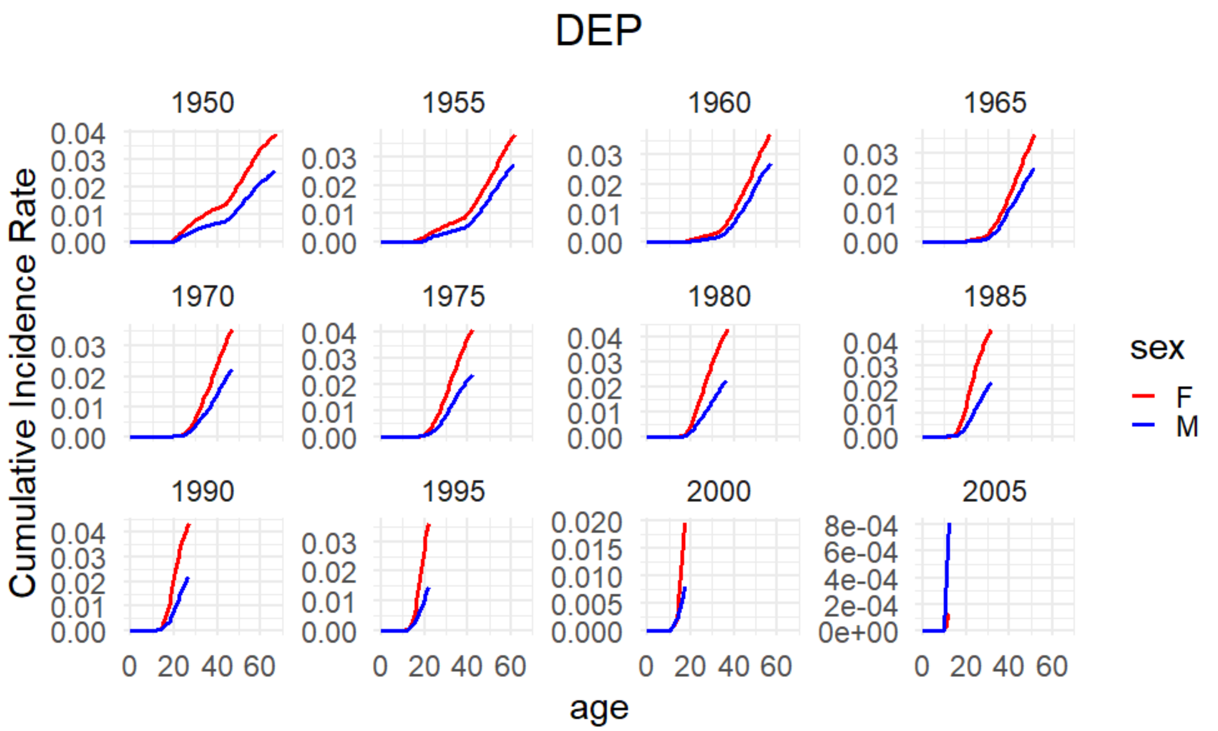


Figure S12: Plot of the cumulative incidence rate for depression grouped by birth year in the Danish registers. The red line corresponds to females and the blue corresponds to males.

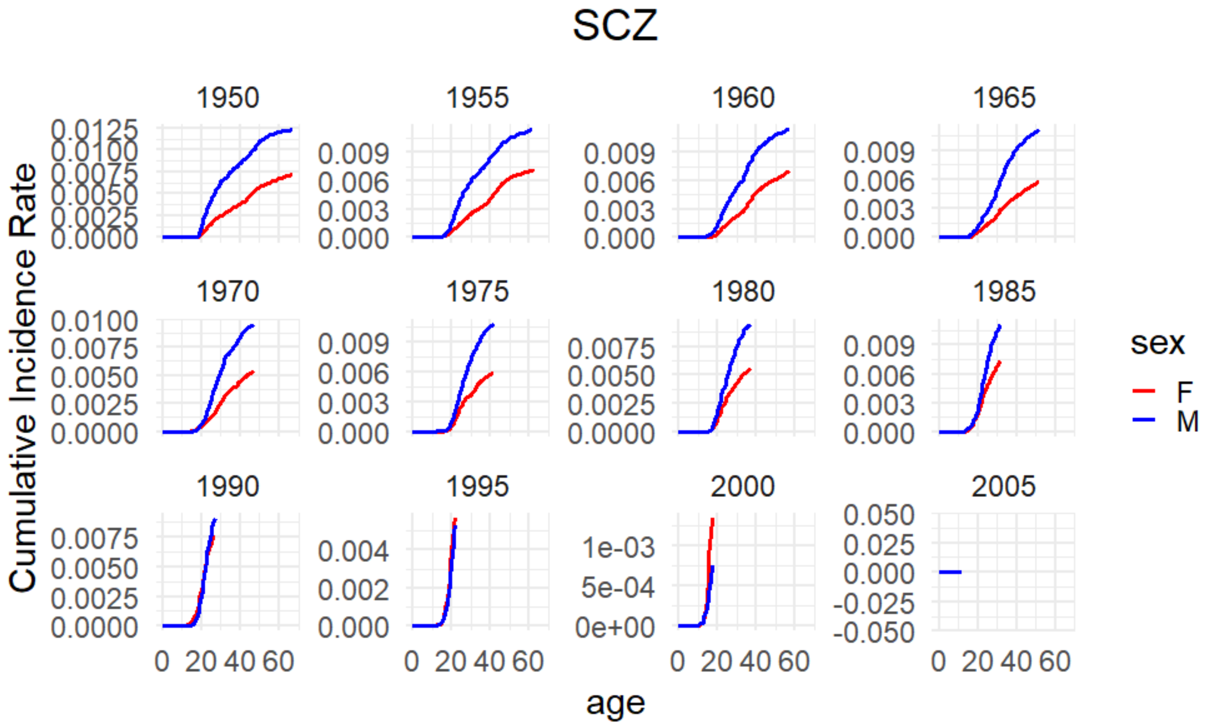


Figure S13: Plot of the cumulative incidence rate for schizophrenia grouped by birth year in the Danish registers. The red line corresponds to females and the blue corresponds to males.

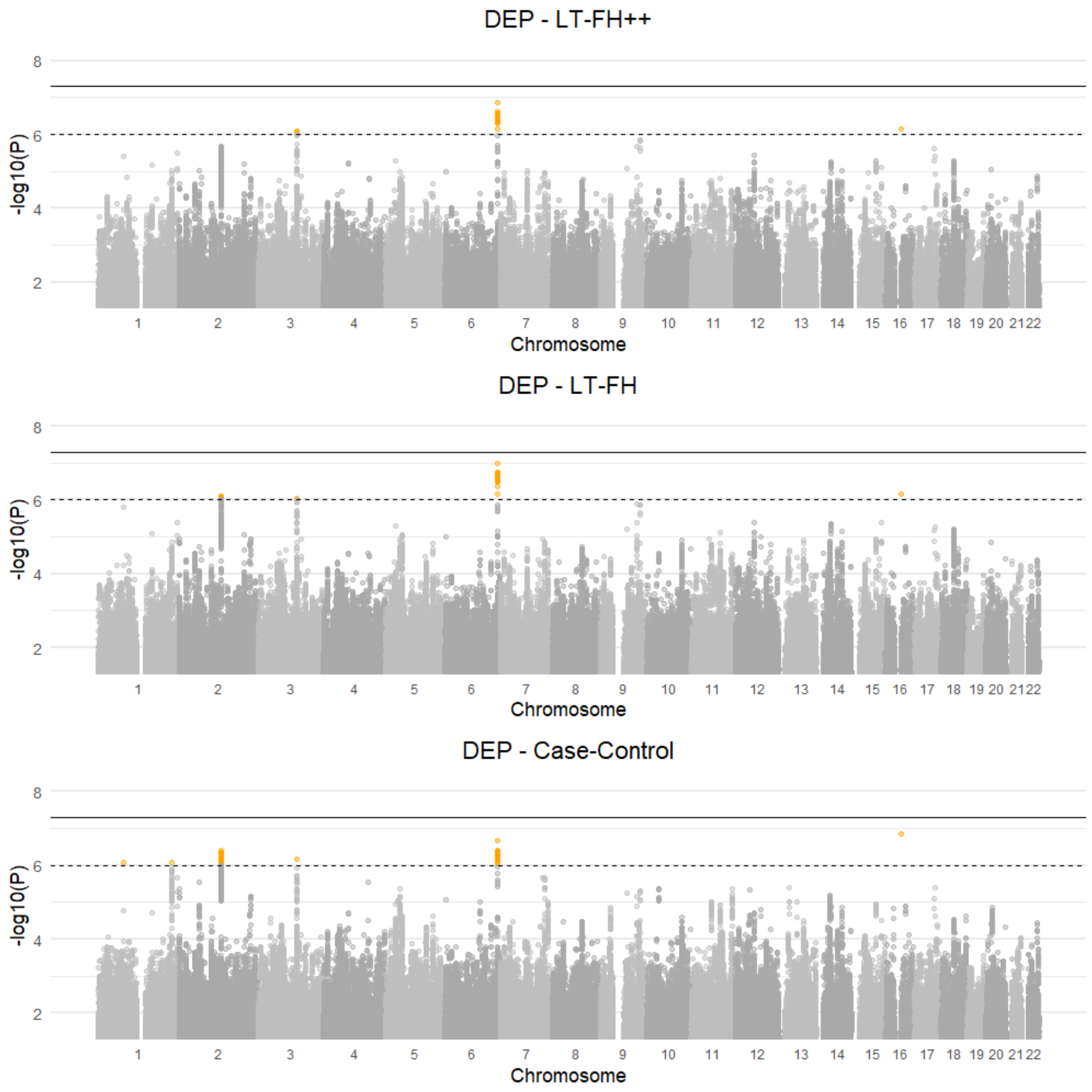


Figure S14: Manhattan plots for LT-FH++, LT-FH, and case-control GWAS of depression in the iPSYCH cohort. The Manhattan plots display a Bonferroni corrected significance level of $5 \times 10^{-8}$, and a suggestive threshold of $5 \times 10^{-6}$. The genome-wide significant SNPs are colored in red and the suggestive SNPs are colored in orange. The squares correspond to top SNPs in a window of size 300k base pairs.

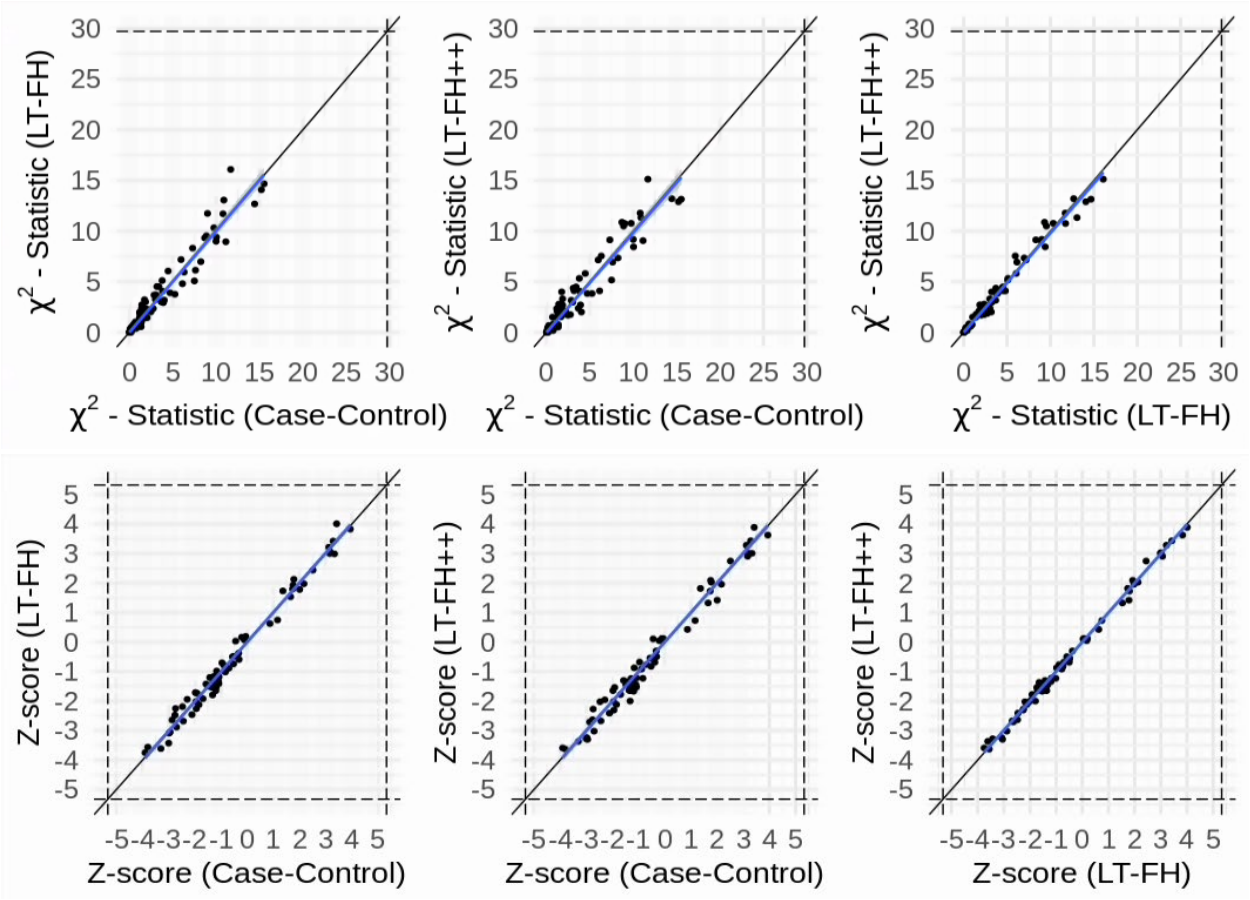


Figure S15: The Z-scores and $\chi^2$ statistics for depression for the three outcomes plotted against each other. The dots correspond to LD clumped SNPs that are genome-wide significant in the largest published meta analysis and present in the iPSYCH cohort (see Methods for details). The blue line indicates the linear regression line between two outcomes and a black line indicates the identity line. The slopes of the regression lines are not significantly different from 1 for any pair of outcomes.

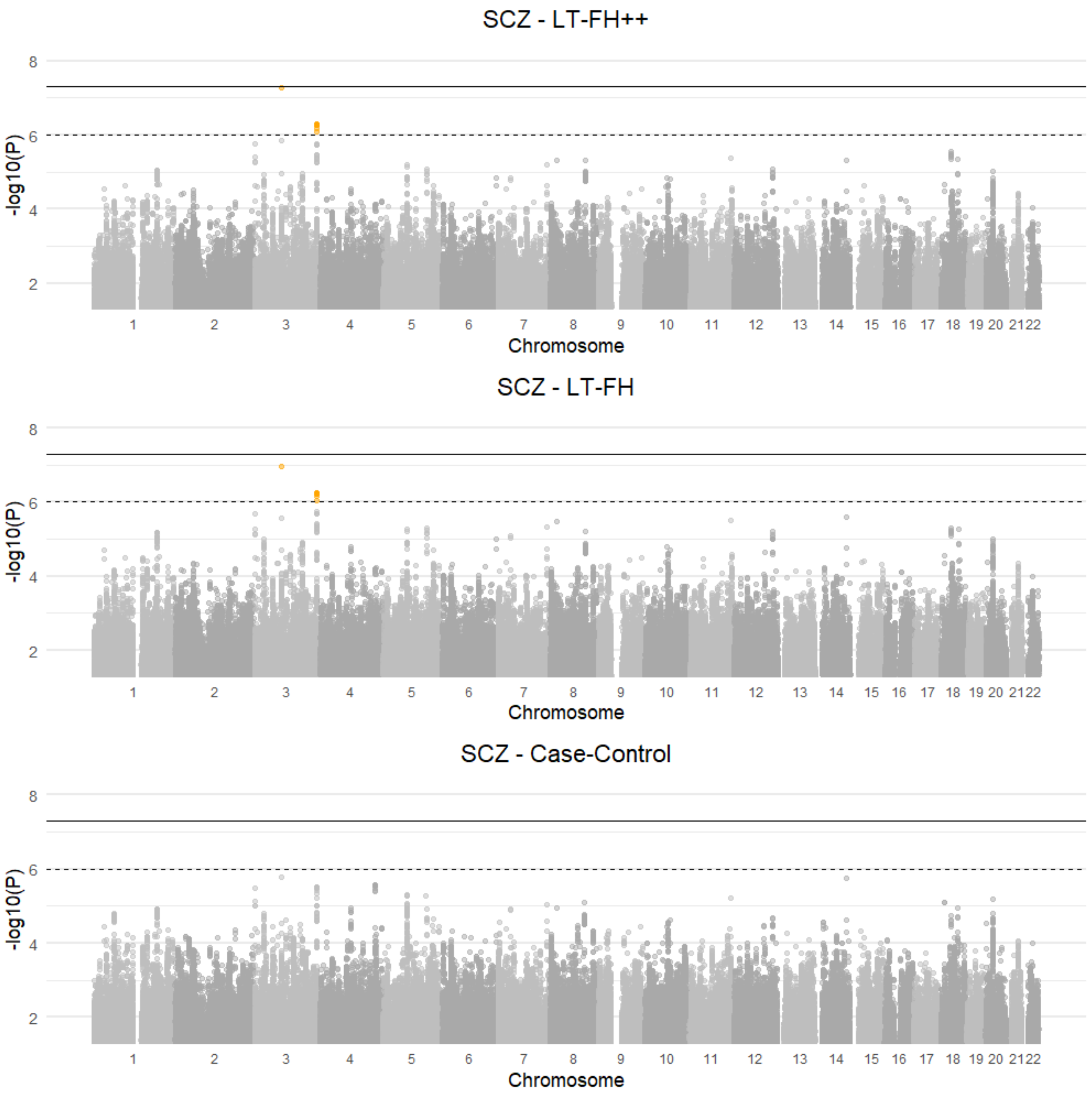


Figure S16: Manhattan plots for LT-FH++, LT-FH, and case-control GWAS of schizophrenia in the iPSYCH cohort. The Manhattan plots display a Bonferroni corrected significance level of $5 \times 10^{-8}$, and a suggestive threshold of $5 \times 10^{-6}$. The genome-wide significant SNPs are colored in red and the suggestive SNPs are colored in orange. The squares correspond to top SNPs in a window of size 300k base pairs.

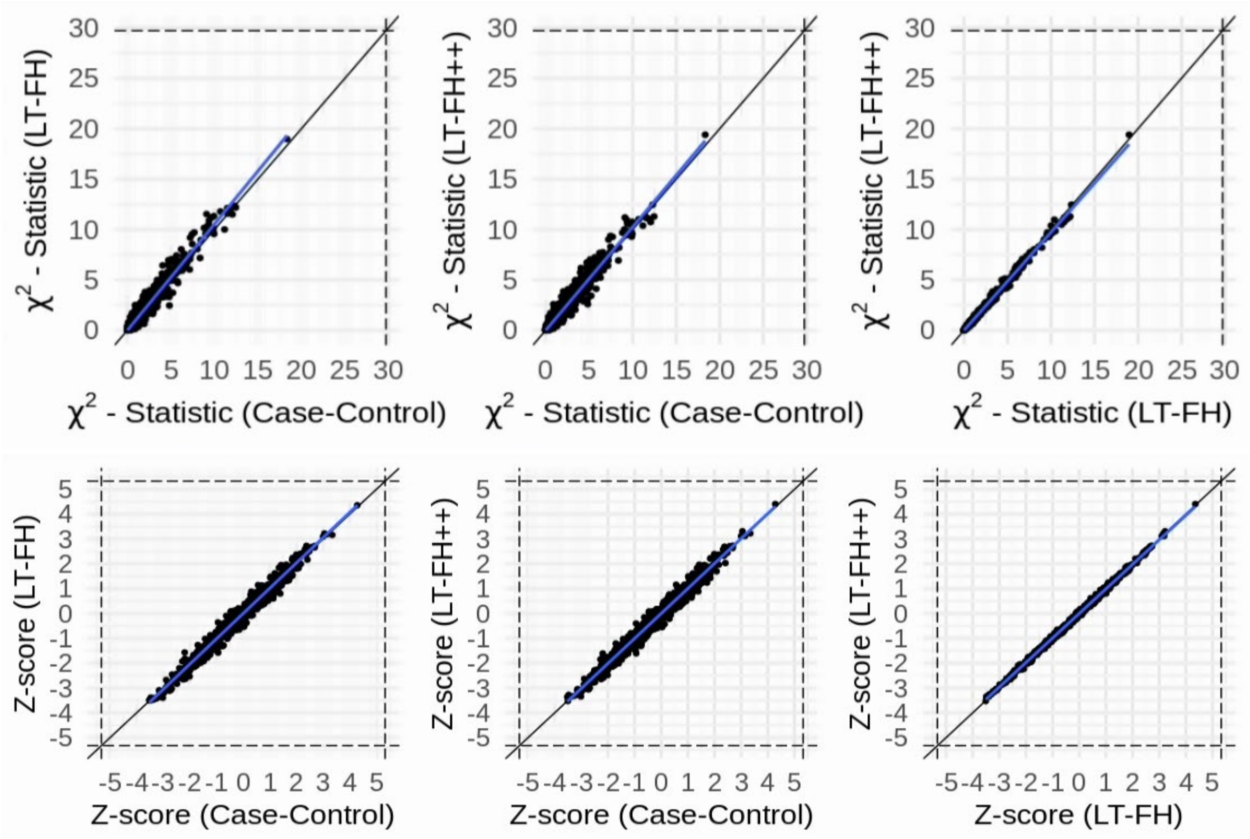


Figure S17: The Z-scores and $\chi^2$ statistics for schizophrenia for the three outcomes plotted against each other. The dots correspond to LD clumped SNPs that are genome-wide significant in the largest published meta analysis and present in the iPSYCH cohort (see Methods for details). The blue line indicates the linear regression line between two outcomes and a black line indicates the identity line. The slopes of the regression lines are not significantly different from 1 for any pair of outcomes.

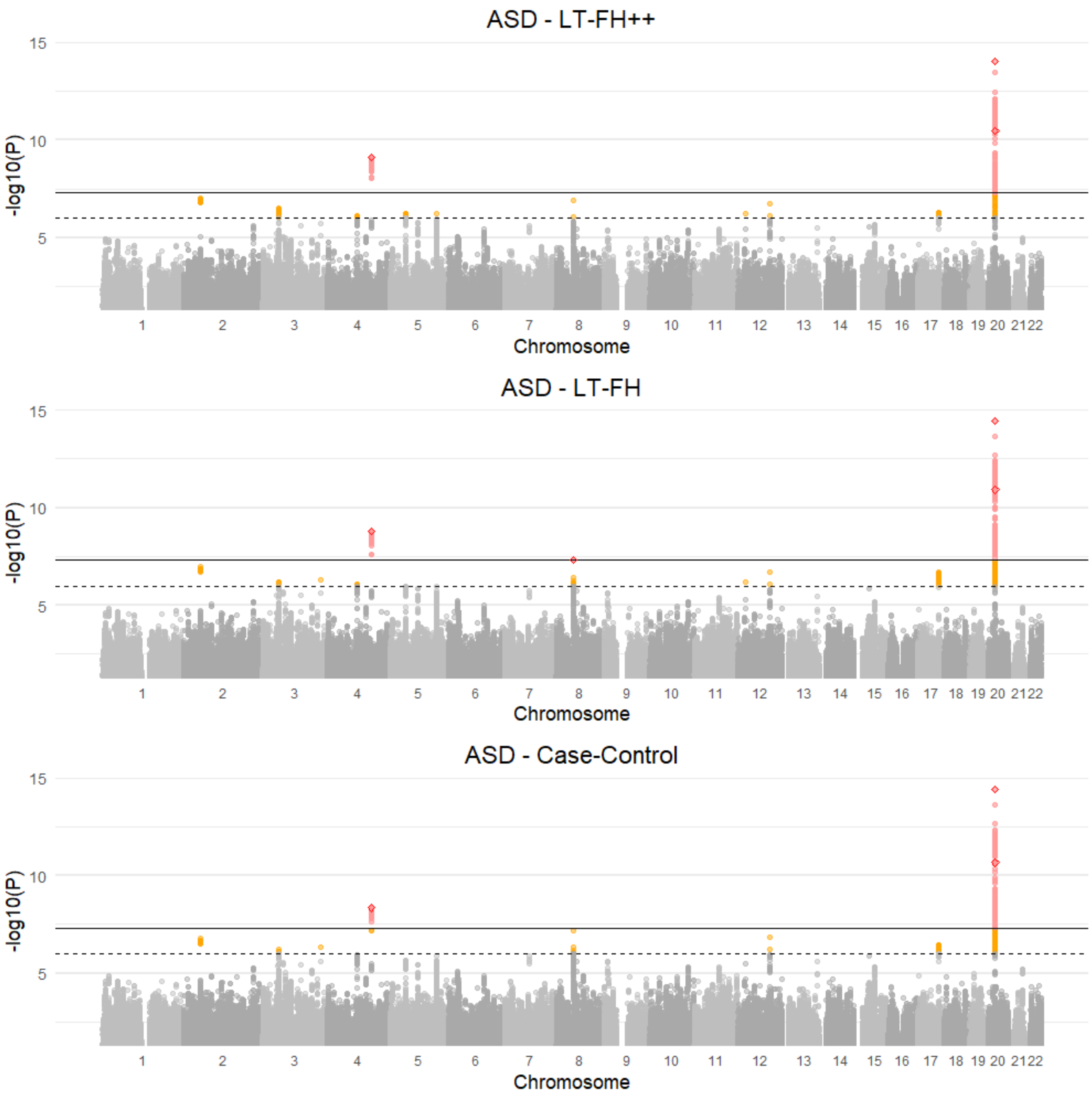


Figure S18: Manhattan plots for LT-FH++, LT-FH, and case-control GWAS of autism spectrum disorder (ASD) in the iPSYCH cohort. The Manhattan plots display a Bonferroni corrected significance level of $5 \times 10^{-8}$, and a suggestive threshold of $5 \times 10^{-6}$. The genome-wide significant SNPs are colored in red and the suggestive SNPs are colored in orange. The squares correspond to top SNPs in a window of size 300k base pairs.

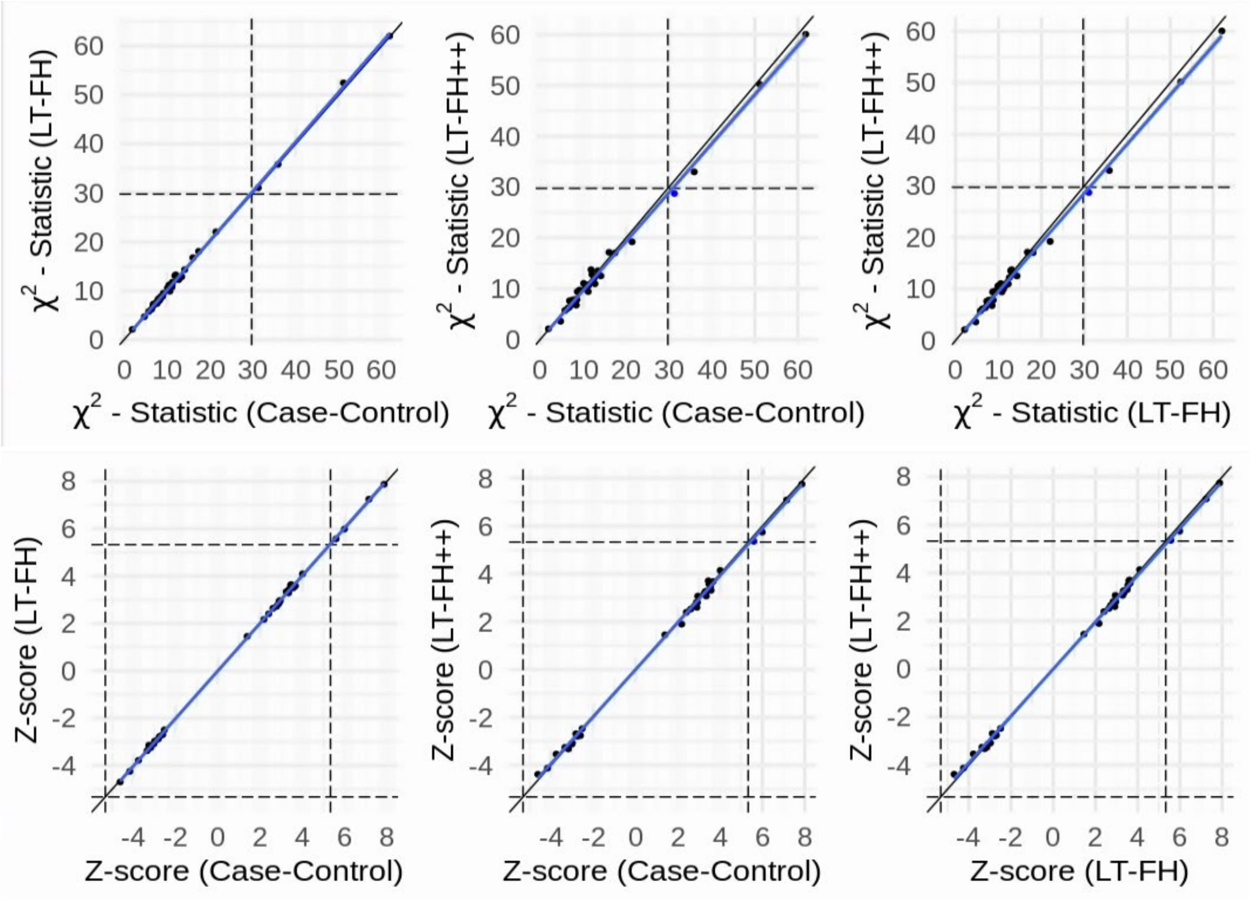


Figure S19: The Z-scores and $\chi^2$ statistics for ASD for the three outcomes plotted against each other. The dots correspond to LD clumped SNPs that are genome-wide significant in the largest published meta analysis and present in the iPSYCH cohort (see Methods for details). The blue line indicates the linear regression line between two outcomes and a black line indicates the identity line. The slopes of the regression lines are not significantly different from 1 for any pair of outcomes.

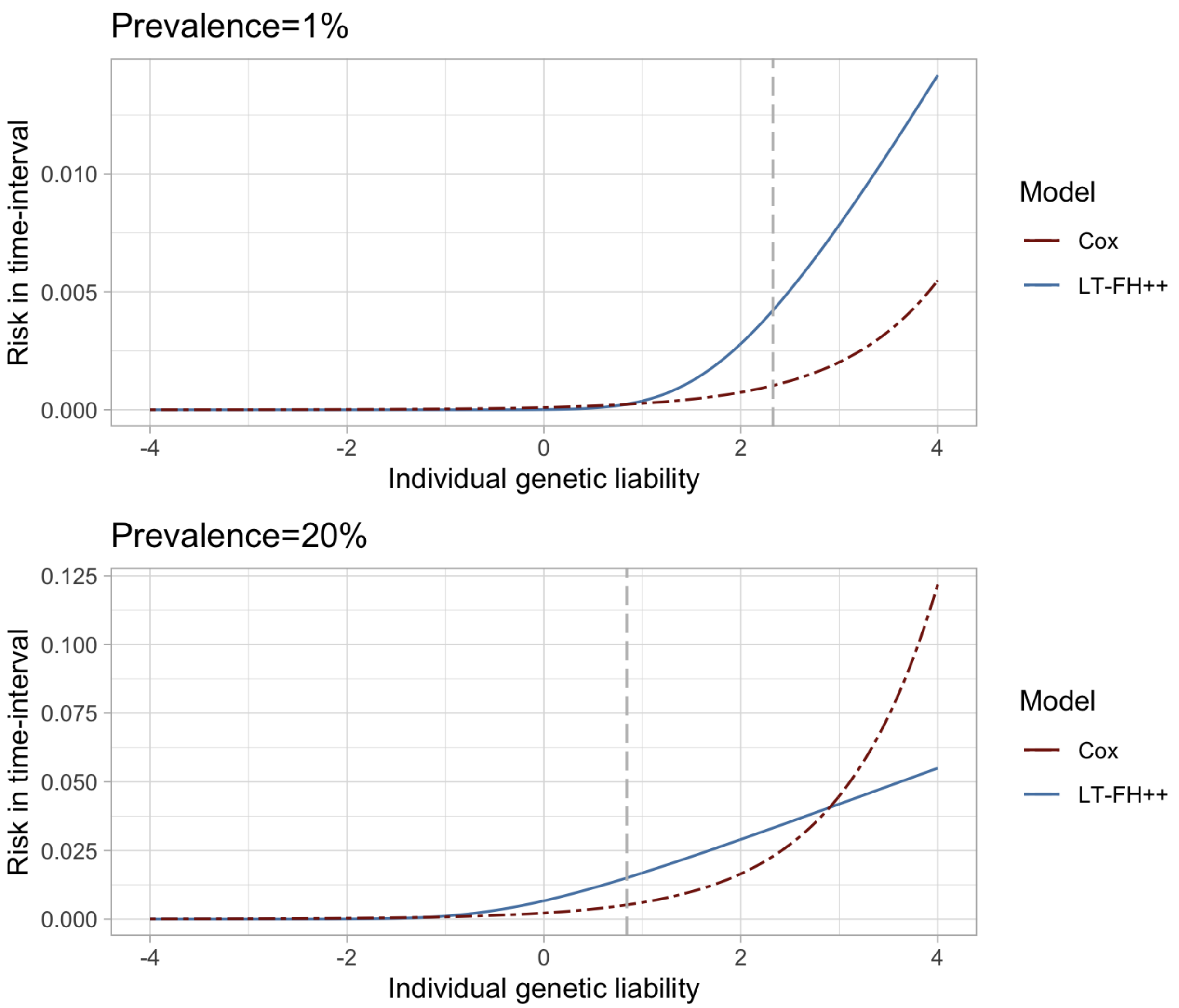


Figure S20: Risk (probability) for becoming a case within a time-interval corresponding to 1% relative increase in prevalence as a function of the genetic liability. The total prevalence changes from 1% and 20% to 1.01% and 20.2% respectively. For Cox regression we assume a constant base incidence rate, corresponding to the prevalence. The vertical dotted grey line denotes the liability threshold corresponding to the prevalence. We note that the risk for becoming a case within a small time-interval is proportional to the hazard rate.

| Variant ID | Chromosome :Position (hg38) | LT-FH++ P-value | Effect size (SE) | Nearest gene | Selected previously reported associations |
|---|---|---|---|---|---|
| rs429358 | 19:44908684 | 8.8e-52 | 0.176493(0.0116573) | APOE | Alzheimer's[7], metabolic traits[76], mortality[42,53] |
| 15:78828640 | 15:78828640 | 1.9e-22 | 0.088522 (0.00908256) | HYKK | Smoking and lung cancer[6], mortality[53] |
| rs10455872 | 6:160589086 | 7.5e-15 | -0.120683 (0.0155212) | LPA | heart disease, mortality[53] |
| 6:161075384 | 6: 161075384 | 5.1e-14 | 0.243674(0.0323606) | MAP3K4 | Endometriosis[77] |
| rs34386495 | 6:32658953 | 4.7e-10 | 0.0664307(0.0106654) | HLA-DQB1 | Asthma[78], autoimmune diseases[79], mortality[53] |
| rs61905747 | 11:113769120 | 8.5e-9 | 0.0620208(0.0107705) | ZW10 | Glioma[80] mortality[53,81] |
| rs2507989 | 6:31356638 | 1.6e-8 | 0.0592997(0.0104863) | HLA-B | White blood cell count[44], Psoriasis[45] |
| rs3838008 | 20:63357289-63357318 (indel) | 1.9e-8 | 0.0608869 (0.0108248) | CHRNA4 | Smoking and lung cancer[6], mortality[53] |
| rs17691989 | 13:77093116 | 4.4e-8 | 0.1571(0.028695) | MYCBP2 | Circadian rhythm (chronotype)[46] |
| rs79339645 | 3:166883110 | 4.7e-8 | 0.120294(0.0220177) | ZBBX | DNA methylation in older people[48] |

**Table S1:** Independent LT-FH++ associations for mortality in UK biobank identified using COJO[43] and sorted by lowest p-value. The two strongest associations are shared with LT-FH, and seven out of three were previously identified in association studies of longevity[81] or parental age[53].

| Variant ID | Chromosome :Position (hg38) | LT-FH++ P-value | Effect size (SE) | Nearest gene | Selected previously reported associations |
|---|---|---|---|---|---|
| rs56022653 | 5:88588020 | 5.8e-12 | 0.132154(0.191985) | LINC00461 | Educational attainment[50], ADHD[10,82] |
| rs11210887 | 1:43610348 | 1.1e-11 | 0.133962(0.0203968) | PTPRF | Smoking initiation[6], Educational attainment[50], ADHD[10,82] |
| rs9969232 | 7:114518899 | 2.1e-9 | 0.120184(0.0200724) | FOXP2 | Risk taking[83], ADHD[10] |
| rs6082363 | 20:21270205 | 5.0e-9 | 0.122019(0.0208684) | ZNF877P | ASD[9,84] |
| rs11030386 | 11:28609701 | 3.7e-8 | 0.106526(0.0193581) | LINC02758 | ADHD[10] |
| rs4261436 | 14:32830276 | 4.3e-8 | 0.103069(0.0188137) | AKAP6 | Cognitive traits[49,50] |
| rs7026534 | 9:134907263 | 4.7e-8 | 0.111291(0.0203778) | | Education attainment, Smoking initiation [6,50] |

**Table S2:** Independent LT-FH++ genome-wide significant associations for ADHD using COJO[43] and sorted by lowest p-value.

| Variant ID | Chromosome :Position (hg38) | LT-FH++ P-value | Effect size (SE) | Nearest gene | Selected previously reported associations |
|---|---|---|---|---|---|
| rs910805 | 20:21248116 | 9.6e-15 | 0.194518 (0.0251149) | ZNF877P, AL117332.1 | ASD[9] |
| rs4274907 | 4:135863730 | 7.7e-10 | 0.173381 (0.0281911) | LOC105377437 | None reported |

**Table S3:** Independent LT-FH++ genome-wide significant associations for ASD using COJO[43] and sorted by lowest p-value.
